# Anaerobic co-digestion of sewage sludge and food waste: staging and carriers enhance system performance and process stability

**DOI:** 10.1101/2025.03.17.643612

**Authors:** Muhammad Ali, Carolina Burgos Pena, Jo De Vrieze

## Abstract

Sewage sludge generated during biological wastewater treatment requires adequate treatment. Anaerobic digestion of sewage sludge is preferred, yet, it is often characterized by low biogas production rates. Co-digestion of sewage sludge with food waste could improve the hydrolysis rate and nutrients balance, resulting in higher biogas production rates. In this study, single- and two-stage anaerobic reactors were operated, with and without carriers in semi-continuous mode for 205 days for assessing the expected improved biogas production rates of sewage sludge and food waste co-digestion. Mono-digestion of sewage sludge resulted in average biogas production rates of 230±31 and 538±40 mL_biogas_ L^-1^ d^-1^ for single- and two-stage systems with carriers, respectively. Co-digestion of sewage sludge with food waste resulted in an improved biogas production rates in both systems. Average biogas production rates of 868±114 and 1507±174 mL_biogas_ L^-1^ d^-1^ were measured in single- and two-stage systems, respectively, with carriers during co-digestion of food waste (30%) and sewage sludge (70%). In all cases, the addition of carriers improved biogas production, and reduced volatile fatty acid accumulation. In conclusion, two-stage co-digestion of sewage sludge and food waste with carriers achieved the highest biogas production and long-term stability.

## 1. Introduction

Rapid population growth, industrialization and depletion of natural resources, along with climate change are obliging humanity to move towards a low carbon, sustainable and resource efficient future [1]. A sustainable bioeconomy can be obtained by converting bio-wastes into valuable, recovered products [2]. This circular approach has engrossed its concern on the municipal solid waste (MSW) and wastewater sectors as key fields that can be improved substantially by implementing waste reduction and waste to energy strategies, like anaerobic digestion [3]. Anaerobic digestion of sewage sludge provides an integrated approach for stabilization of organic matter, pathogens removal, nutrients, and energy recovery [4,5]. This process is also characterized by slow hydrolysis rates, and, thus, requires large bioreactors for sludge treatment [4,6–11]. Anaerobic digestion of food waste is distinguished with rapid hydrolysis and acidification, due to accumulation of volatile fatty acids (VFA), and can result in process failure, if not controlled well [6,12–15]. Mono-digestion of food waste and several other feedstocks is challenging, mainly due to nutrients imbalance and lack of diverse microbial community, which may result in lower biogas yields [8].

Co-digestion is a common strategy that could enable higher biogas yields, balanced nutrients availability, toxicity dilution, and can potentially overcome limitations associated with mono-digestion of sewage sludge and food waste [3,8,10,13,16,17]. It can also offer enhanced system stability and methane yields through synergetic effects of a diverse microbial community and improved buffering capacity [3,8,18,19].

Two-stage anaerobic digestion is a promising way to further improve (co-)digestion efficiency, by separating the different stages of microbial activity, resulting in better process control compared to the single-stage systems. Many researchers have investigated the two-stage systems for improving system performance and stability treating various feedstocks, like sewage sludge, food waste and animal manure, in comparison with single-stage systems [3,20–24]. The first-stage acts as a pretreatment step in which fermentative bacteria make substrate accessible to the methanogenic archaea in the second stage, thus, resulting in improved methane production [3,25–28]. The first-stage is usually operated at a short hydraulic retention time (HRT) of 1 to 3 days, and second-stage at long HRT of 14 to 40 days for providing favourable conditions to fermentative bacteria and methanogenic archaea, respectively [29–31]. Higher biogas and methane yields and volatile solids (VS) removal efficiencies of 35%, 60% and 9%, respectively, have been reported in two-stage anaerobic co-digestion, compared with single-stage anaerobic co-digestion of sewage sludge and food waste [20,25,27,29,31]. In other studies, two-stage systems showed superiority in terms of biogas quality and organics removal at organic loading rates (OLR) up to 5 g_VS_ L^-1^ d^-1^, compared with single-stage systems treating sewage sludge and food waste [33–35]. Two-stage systems overall produce more biogas, and are more effective in overcoming difficulties, such as pH drops, due to volatile fatty acids (VFA) accumulation, which ceases biogas production in a single-stage system [35].

Despite its benefits, co-digestion is a complex process, with stability and optimization issues that even two-stage systems cannot overcome [8]. The use of carriers in the digestion process might assist in process stability, as different studies have shown the effectiveness of carriers in co-digestion process [37–40]. Carriers provide an additional surface area to slow growing methanogens archaea, which results in better system performance by keeping the methanogenic community at optimal conditions in the digester. This attributes to an improved VFA assimilation, resulting in less residual VFA, and, thus, overall better system stability, compared to systems without carriers [40].

In this study, a two-stage system was designed with an intent to increase overall biogas production by (1) producing biogas from both the stages and (2) retaining the methanogenic community through the addition of carriers in the single-stage system and in stage-II of the two-stage system. The key objectives of the study were: (1) to maximize the overall biogas production in two-stage system co-digesting sewage sludge and food waste by operating the first-stage at a HRT (5 days) to produce more biogas instead of bio-hydrogen by promoting hydrogenotrophic methanogens activity; (2) to increase biogas yields, through the effectiveness of carriers usage to enrich slow growing methanogens; and (3) to compare the overall system performance of single- and two-stage systems (with and without carriers) in terms of biogas production, residual VFA accumulation and VS reduction.

## 2. Material and methods

### 2.1 Inoculum and feedstocks

The inoculum and sewage sludge used for all experiments were collected from a local wastewater treatment plant with an anaerobic digester, operated by Aquafin, and located in Ghent, Belgium. Food waste was collected from the canteen of the Faculty of Bioscience Engineering, Ghent University, which mostly comprised of vegetables, rice, meat, and bread. Food waste was diluted 10% with tap water by adding 10 g of water to 90 g of food waste, homogenized by using a kitchen blender for making a slurry, and stored at 4°C before use. The main characteristics of the inoculum and sewage sludge and food waste (co-)feedstock are presented in Table 1.

**Table 1.** Characterization of the inoculum and feedstocks used in the different experiments. All analyses were conducted in analytical triplicates. Data are expressed as mean ± standard deviation of the analytical triplicates. ND = not determined.

| Parameter | Unit | Inoculum | Thickened<br>Sludge (T.<br>S) | Sewage<br>Sludge (S. S) | Food Waste<br>(FW) |
| --- | --- | --- | --- | --- | --- |
| pH | - | 8.25±0.05 | 7.21±0.11 | 6.39±0.04 | 5.93 ± 0.07 |
| Electrical<br>Conductivity (EC) | mS cm<sup>-1</sup> | 7.79±0.03 | 18.09±1.34 | 3.96±0.02 | ND |
| Total Solids (TS) | g L<sup>-1</sup> | 39.2±2.7 | 73.0±2.4 | 8.6±0.5 | 134±11 |
| Volatile Solids<br>(VS) | g L<sup>-1</sup> | 22.0±2.1 | 47.8±1.7 | 5.4 ± 0.4 | 124±9 |
| Volatile solids (%<br>of TS) | % | 56.1±0.3 | 65.4±0.0 | 62.8 ± 1.1 | 92.5±0.1 |
| Total chemical<br>oxygen demand<br>(TCOD) | g L<sup>-1</sup> | 30.3±8.1 | 79.3±22.9 | 3.7± 0.2 | 195±17 |
| Soluble chemical<br>oxygen demand<br>(SCOD) | g L<sup>-1</sup> | 1.60±0.22 | 7.01±2.52 | 0.06 ± 0.01 | 48±4 |
| Total Nitrogen<br>(TN) | g L<sup>-1</sup> | 3.80±0.68 | 2.71±0.30 | 0.64±0.11 | 5.73±0.52 |
| Carbon to<br>nitrogen ratio<br>(COD/N) | - | 7.9±0.8 | 29.3±0.1 | 5.7±0.3 | 34.0±0.0 |

### 2.2 Reactor set-up and operation

Four different treatments, *i.e.*, a single stage and two-stage system, both with and without carriers, were performed in biological duplicates using sewage sludge and food waste as main (co-)feedstock. All treatments were operated in glass Schott bottles (500 mL) that were used as reactors with a working volume of 400 mL. Reactors were placed in a water bath for maintaining a constant mesophilic temperature (35°C), and were connected with plastic cylinders that were used as gas columns for biogas quantification through a water displacement method (Figure S1). The initial inoculum biomass concentration of 10 g_VS_ L^-1^ (volatile solids) was fixed by diluting the inoculum with tap water.

During the acclimatization phase, reactors were operated by feeding thickened sewage sludge for 8 days, initially at a HRT of 80 days, and an OLR of 0.6 g_VS_ L^-1^ d^-1^ was applied to allow adaptation of microbial community to the incoming feedstock. Next, the reactors were operated for 13 and 17 days at a HRT of 40 and 20 days, respectively. The period after the acclimatization phase (in total 38 days) served as the experimental period, and lasted for 205 days.

High-density polyethylene carriers with a specific surface area of 800 m² m^-3^ (L9.1mm, Anox Kaldness K5, Lund, Sweden) with a filling ratio of 20% were used in the single- and two-stage systems. Both the single- and two-stage treatments (with and without carriers) were operated at a total HRT of 20 days. In stage-I and stage-II (with and without carriers) of the two-stage system, a HRT of 5 and 15 days was fixed, respectively. By increasing the food waste proportion (from 10% to 40% on wet mass basis till the reactors crashed) in the co-feedstock during co-digestion of sewage sludge and food waste, the OLR was stepwise increased (Table 2).

**Table 2.**
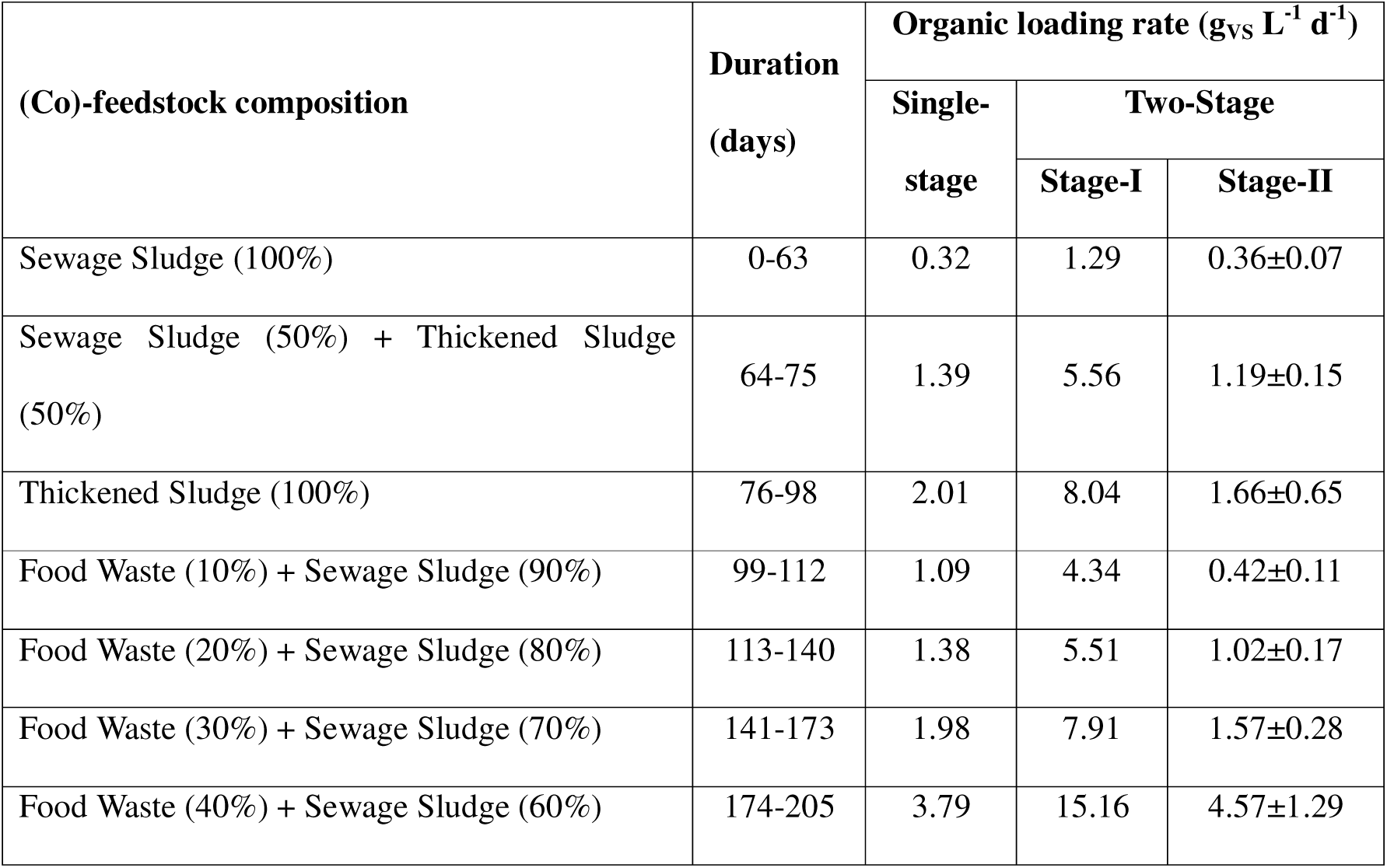
The different phases of the single-stage and two-stage co-digestion reactors (with and without carriers) operated at varying organic loading rates (OLRs) and different (co-)feedstock composition.

| (Co)-feedstock composition | Duration<br>(days) | Organic loading rate ($\text{g}_{\text{VS}} \text{L}^{-1} \text{d}^{-1}$) | | |
| --- | --- | --- | --- | --- |
|  |  | Single-stage | Two-Stage |  |
|  |  |  | Stage-I | Stage-II |
| Sewage Sludge (100%) | 0-63 | 0.32 | 1.29 | 0.36±0.07 |
| Sewage Sludge (50%) + Thickened Sludge (50%) | 64-75 | 1.39 | 5.56 | 1.19±0.15 |
| Thickened Sludge (100%) | 76-98 | 2.01 | 8.04 | 1.66±0.65 |
| Food Waste (10%) + Sewage Sludge (90%) | 99-112 | 1.09 | 4.34 | 0.42±0.11 |
| Food Waste (20%) + Sewage Sludge (80%) | 113-140 | 1.38 | 5.51 | 1.02±0.17 |
| Food Waste (30%) + Sewage Sludge (70%) | 141-173 | 1.98 | 7.91 | 1.57±0.28 |
| Food Waste (40%) + Sewage Sludge (60%) | 174-205 | 3.79 | 15.16 | 4.57±1.29 |

Reactors were operated in a semi-continuous mode by feeding them manually three times per week by replacing digestate with the fresh (co-)feedstock in single-stage system to maintain the total HRT of 20 days. For the two-stage system, fresh (co-)feedstock replaced the digestate in stage-I and stage-II was fed by replacing its digestate with the digestate of stage-I to maintain the desired total HRT of 5 and 15 days in stage-I and stage-II of two-stage system, respectively, during the entire experimental period. Biogas production and composition were also monitored thrice a week. The results of biogas and methane production were reported at standard temperature (273.15 K) and pressure (101.325 kPa) (STP) conditions. Digestate samples were collected before each feeding (three times per week), and stored at 4°C until use for analytical testing on weekly basis. The pH was measured three times per week, but was not corrected.

### 2.3 Analytical techniques

The pH was measured with a C532 pH meter, and conductivity was measured with a C833 electrical conductivity (EC) meter (Consort, Turnhout, Belgium). Total solids (TS) and volatile solids (VS) were determined according to the Standard Methods (APHA, 2005). Total nitrogen (TN) and the total chemical oxygen demand (TCOD) were measured by using nanocolor tube tests (Hach LCK 514, Hach-Lange, Germany). For soluble chemical oxygen demand (sCOD) measurements, samples were prefiltered through 0.45 µm polyamide filters (Hach LCK 514, Hach-Lange, Germany). These test tubes were digested using a nanocolor Vario 4 (Macherey-Nagel, Germany), and the COD concentration was measured using a nanocolor 500 D spectrophotometer (Macherey-Nagel, Germany). Volatile fatty acids (VFA) were determined by means of capillary gas chromatography (GC-214), coupled with flame ionization detector (FID) with autosampler (Shimadzu®, The Netherlands) with a DB-FFAP 123–3232 column (30 m x 0.32 mm × 0.25 µm; Agilent, Belgium) and a flame ionization detector (FID), calibrated for VFA concentration range of 30–1000 mg/L using a nitrogen gas carrier. The COD-adjusted VFA values were calculated by multiplying the measured acid concentration by the ratio of the oxygen required for oxidation to CO_2_ to the molecular weight of the acid. The biogas composition (CH_4_, H_2_ and CO_2_) was analysed with a compact gas chromatograph (Global Analyser Solutions, Breda, The Netherlands). Two channels measured CH_4_, O_2_, N_2_ and H_2_ (Porabond Q pre-column and Molsieve 5A column), and CO_2_ (Rt-QS-bond column and pre-column), respectively. The detection limit was 100 ppmv through thermal conductivity detector. Anion and cation concentrations were measured using ion chromatography (Metrohm, Switzerland) using a Metrosep A Supp 5–150/4.0 (61006520) column. Detection limits for ions ranged between 0.05 and 100 mg L^-1^. The Origin (version pro-2021), and Microsoft Excel were used for data analysis and fitting.

## Results

### 2.4 Biogas production and methane yield

During the first 62 days of operation after the start-up phase, a biogas production rate of 194±28 and 487±52 mL_bioags_ L^-1^ d^-1^ was observed in the single- and two-stage systems, respectively, without carriers treating 100% sewage sludge (Figure 1). An average biogas production rate of 230±31 and 538±40 mL_bioags_ L^-1^ d^-1^ was measured in the single- and two-stage systems, respectively, with carriers, treating the same feedstock. The methane yield in the single- and two-stage systems without carriers during the first 62 days of operation when treating 100% sewage sludge ranged between values of 171±56 and 301±49 mL CH_4_ g^-1^_vs_ respectively. With an increase in the OLR from 1.2 to 8 g_VS_ L^-1^ d^-1^, by replacing sewage sludge with thickened sludge, a substantial increase in biogas production rate and rapid decline in methane yield was noticed from days 63 to 97 (Figure 2a). In the two-stage system, the stage-I reactor showed a consistent low methane yield under 200 mL CH_4_ g^-1^_vs_, and the stage-II reactor exhibited a decreasing methane yield prior to the introduction of food waste until day 98 (Figure 2b).

**Figure 1:**
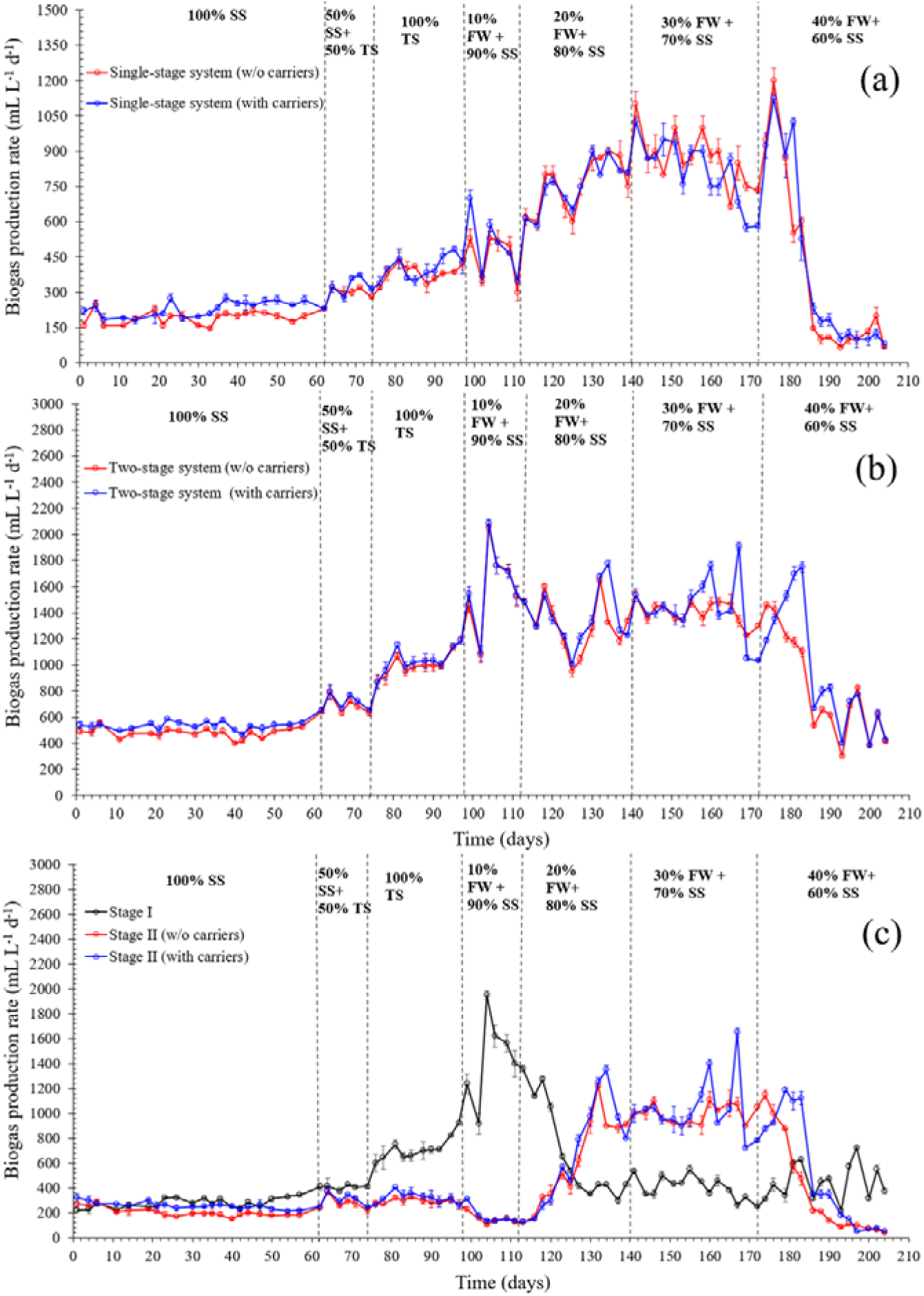
Biogas production rate in (a) the single-stage anaerobic co-digestion (SS-AncoD) system without carriers (red line) and with carriers (blue line) during the entire operational period (205 days) excluding start-up time; (b) two-stage anaerobic co-digestion (TS-AncoD) system without carriers (red line) and with carriers (blue line); (c) the stage-I (black line), stage-II without carriers (red line) and stage-II with carriers (blue line) of the two-stage system. The dashed, black line indicates the beginning of the new feeding regimes. Data is expressed as mean ± standard deviation of biological duplicates (n=2). SS = Sewage Sludge, TS = Thickened Sludge, and FW = Food Waste.

**Figure 2:**
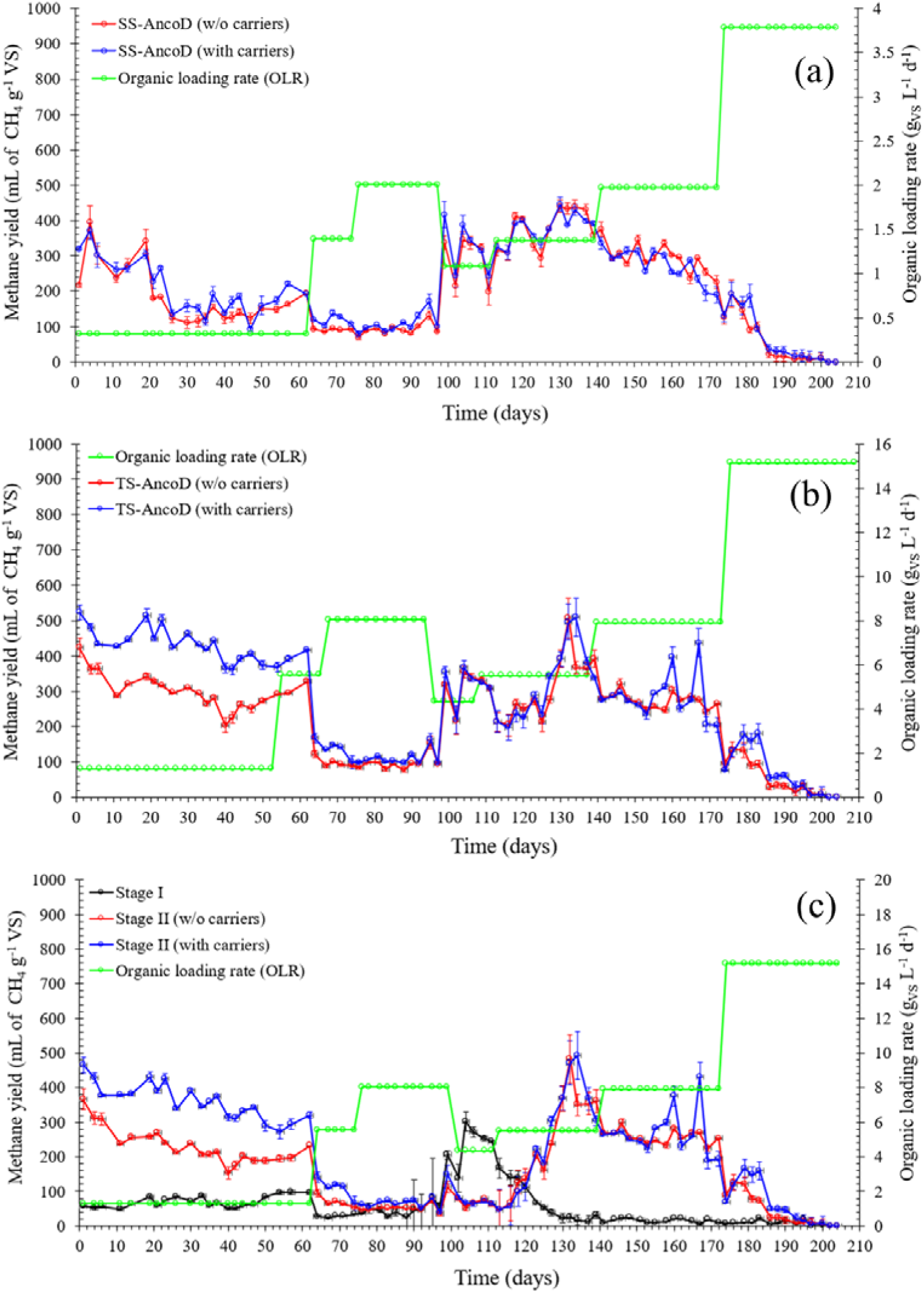
Methane yield of (a) single-stage anaerobic co-digestion (SS-AncoD) system without carriers (red line) and with carriers (blue line), (b) two-stage anaerobic co-digestion (TS-AncoD) system without carriers (red line) and with carriers (blue line), and (c) stage-I (black line), stage-II without carriers (red line) and stage-II with carriers (blue line) of two-stage system. The organic loading rate is presented by the green line. Data is expressed as mean ± standard deviation of biological duplicates (n=2).

In the next phase, when food waste was added as co-feedstock, biogas production rate increased to 696±133 and 1922±403 mL_biogas_ L^-1^ d^-1^, respectively, in the single- and two-stage systems with carriers. An increase in the methane yield was observed in stage-I and stage-II reactors of two-stage system from days 99 to 112, linked to the enhanced biogas production rate during this period (Figure 1b). The methane yield was higher in stage-II reactors, due to lower organic loading rate and higher average methane content of 87±3% in comparison with a methane content of 75±4% in the stage-I reactors (Figure S2) treating 10% food waste and 90% sewage sludge co-feedstock. A temporal decline in biogas production was observed in both reactors when the food waste fraction was progressively increased to 20% in the co-feedstock. The methane yield decreased to values below 50 mL CH_4_ g^-1^_vs_ in the stage-I reactor from day 127 onwards, while the stage-II reactor with carriers reached a maximum methane yield of 490±24 mL CH_4_ g^-1^_vs_ on day 134 at an OLR of 1.0 g_vs_ L^-1^ d^-1^. When the food waste fraction was further increased to 30%, a maximum methane yield of 302±51 and 425±34 mL CH_4_ g^-1^_vs_ were observed on day 146 and 167 in stage-II reactors of the two-stage system without and with carriers, respectively. A further increase in food waste up to 40% in the co-feedstock did not result in more biogas, but a sharp decline in biogas production was noticed in both reactors after day 202 until the end of experimentation period. A downward trend of methane yield was also observed, with values decreasing below 50 mL CH_4_ g^-1^_vs_ in the stage-II reactors (without and with carriers) on day 183 and 190, respectively. Overall, in both single- and two-stage systems, the use of carriers had a positive impact on the methane content, and the two-stage system achieved a higher biogas production rate and methane yield than the single-stage reactor treating different (co)-feedstocks.

### 2.5 pH and volatile fatty acids

The pH is considered a critical operating parameter in the anaerobic digestion process, because acidification is one of the major reason of reactors failure, due to VFA accumulation, which results in a pH drop. The pH values in the single-stage reactors, without and with carriers, were in the stable range of 7.2 - 7.9 till day 176 at the start of 40% food waste and 60% sewage sludge co-feedstock regime (Figure 3). From day 179 till the end of the experiment, the pH decreased rapidly in both single-stage reactors without and with carriers to values of 5.4±0.2 and 5.4±0.0 on day 186 and 197, respectively, indicating a rapid acidification within the reactors at a maximum OLR of 3.8 g_VS_ L^-1^ d^-1^, which was reflected in a decrease in biogas production rates, as discussed earlier. This pH decline was more pronounced in the reactor without carriers, suggesting a slightly improved system stability in the reactors with carriers.

**Figure 3:**
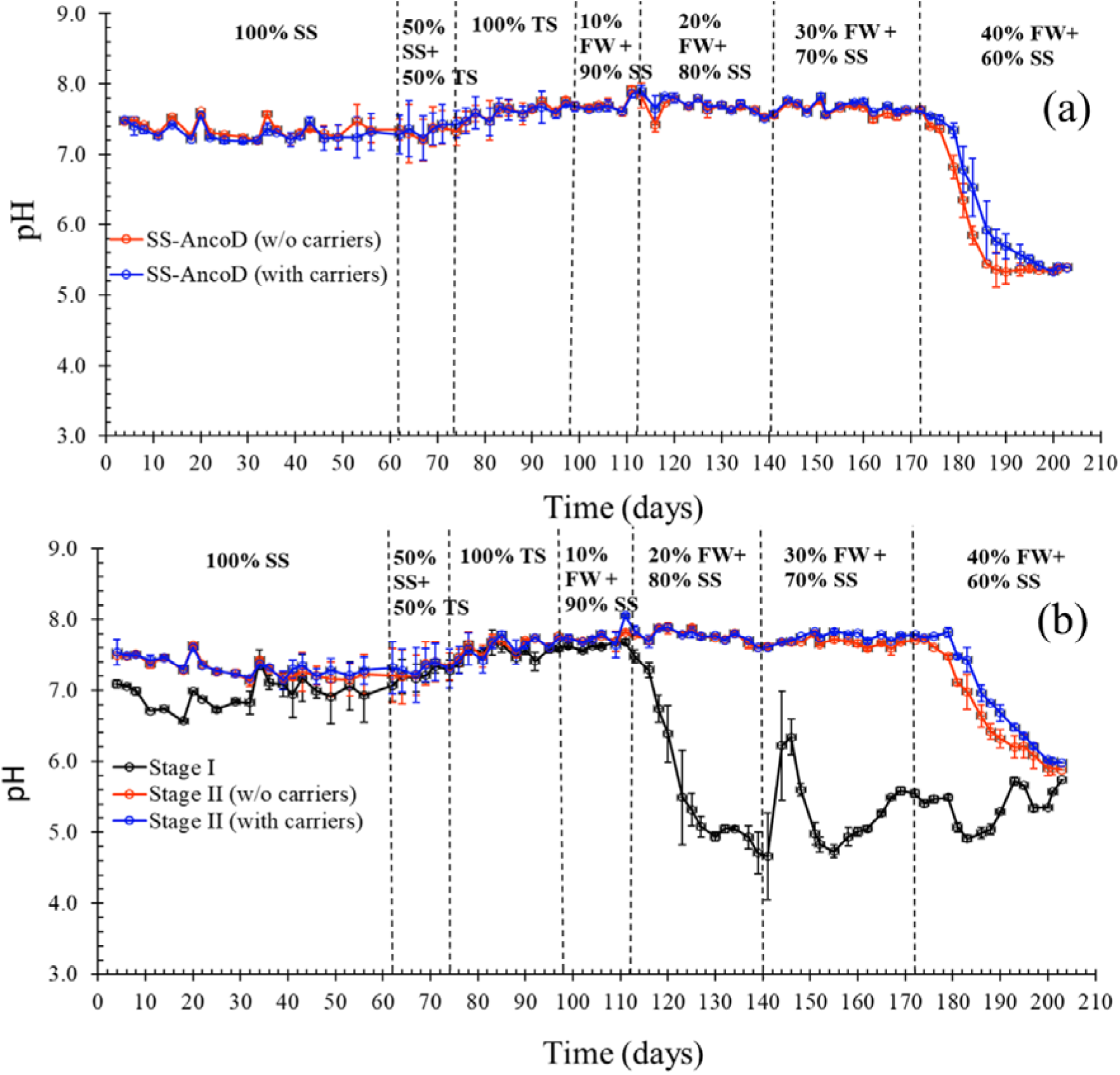
The pH in (a) the single-stage anaerobic co-digestion (SS-AncoD) system without carriers (red line) and with carriers (blue line), (b) stage-I (black line), stage-II without carriers (red line) and stage-II with carriers (blue line) of the two-stage system. Data is expressed as mean ± standard deviation of biological duplicates (n=2). SS = Sewage Sludge, TS = Thickened Sludge, and FW = Food Waste.

In the two-stage systems, a decline in pH value in stage-I was observed on day 120 at an OLR of 5.5 g_VS_ L^-1^ d^-1^, which refers to 20% food waste and 80% sewage sludge co-feedstock. The addition of 20% food waste caused a fluctuation in pH levels in stage-I (Figure 3b). The pH in stage-I reactor dropped to a minimum value of 4.7±0.8 at the start of the 30% food waste and 70% sewage sludge application on day 141, and increased again to a value of 5.7±0.0 at the end of the experiment, referring to highest OLR of 15.2 g_VS_ L^-1^ d^-1^. A substantial drop in pH from 7.6±0.0 to 5.9±0.1 and 7.6±0.0 to 6.0±0.0 in stage-II reactors without and with carriers, respectively, was observed on day 203. Similar to the single-stage system, the pH drop occurred earlier in the two-stage reactors without carriers than in the reactors with carriers (Figure 3b).

A total VFA concentration of 0.2±0.0 g_COD_ L^-1^ was measured during mono-digestion of sewage sludge, and it slowly increased to 1.5±0.7 g_COD_ L^-1^ and 0.6±0.1 g_COD_ L^-1^ during co-digestion of 30% food waste and 70% sewage sludge in the single-stage system without and with carriers, respectively. When the food waste fraction was increased from 30 to 40%, an increase in the VFA accumulation was also observed, with total VFA levels reaching up to 11.4±4.0 and 18.7±1.1 g_COD_ L^-1^ in the single-stage reactors with and without carriers, respectively (Figure 4a), which caused the pH drop in the reactors (Figure 3a), as discussed earlier.

**Figure 4:**
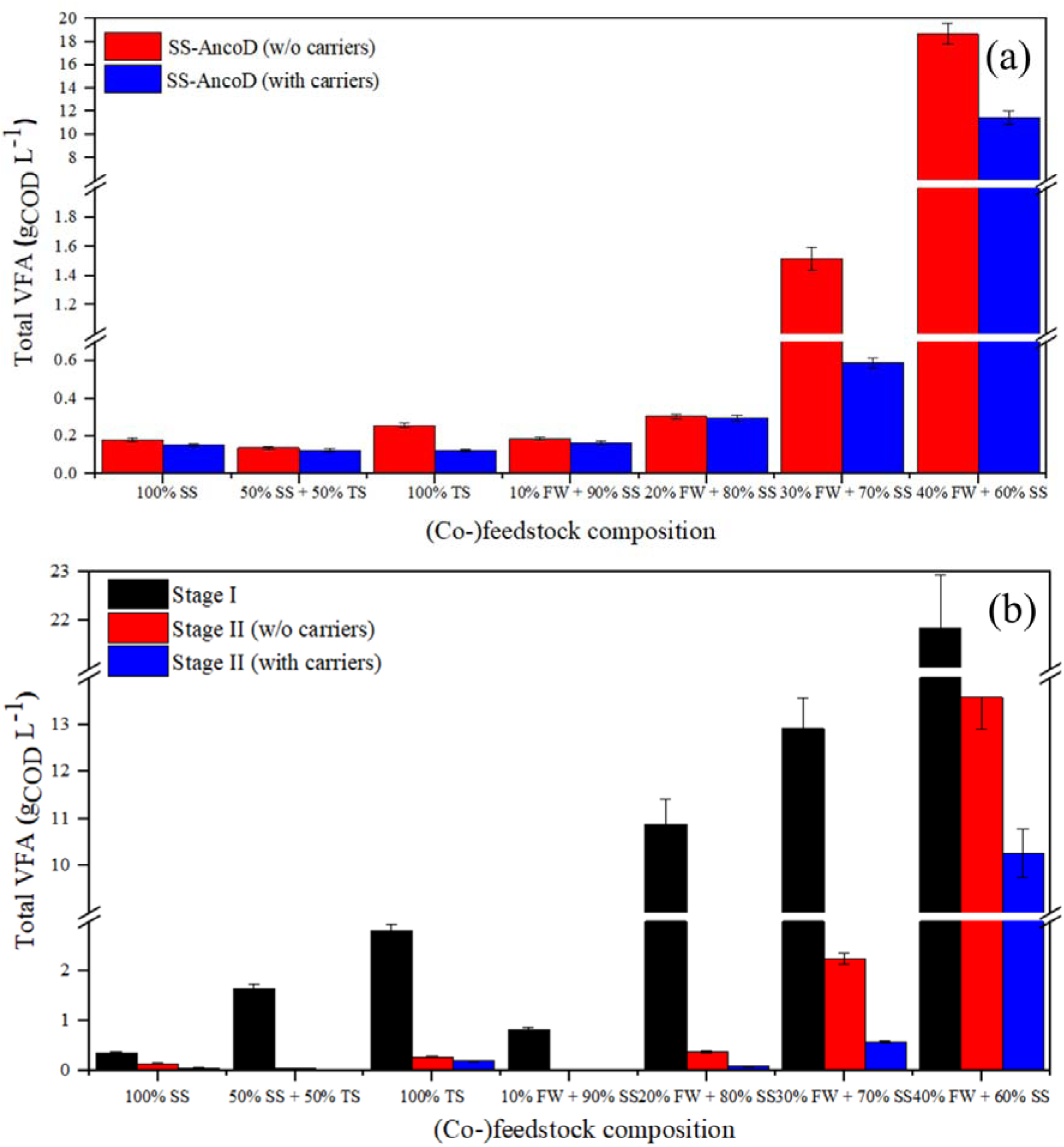
Total volatile fatty acid (VFA) in (a) the single-stage anaerobic co-digestion (SS-AncoD) reactor without carriers (red block) and with carriers (blue block), and in (b) stage-I (black block), stage-II without carriers (red block) and stage-II with carriers (blue block) of the two-stage system. Data is expressed as mean ± standard deviation of biological duplicates (n=2). SS = Sewage Sludge, TS = Thickened Sludge, FW = Food Waste, VFA = Volatile Fatty Acids, and COD = Chemical Oxygen Demand.

Stage-I of the two-stage system showed an increase in VFA concentration from 0.4±0.0 to 2.8±0.2 g_COD_ L^-1^ by changing the feedstock from 100% sewage sludge to 100% thickened sludge. With the gradual addition of food waste in to the feedstock, an increase in VFA concentration from 0.8±0.1 to 21.8±1.8 g_COD_ L^-1^ was measured throughout the experiment, due to an increased OLR from 4.3 to 15.2 g_VS_ L^-1^ d^-1^. The stage-II reactors with carriers showed less VFA accumulation in comparison with the reactors without carriers, which indicates higher reactor stability. A total VFA concentration of 0.14±0.00 g_COD_ L^-1^ was measured in stage-II of two-stage systems without carriers treating 100% sewage sludge, and this increased to 13.57±7.78 g_COD_ L^-1^ with 40% food waste and 60% sewage sludge as co-feedstock. A VFA concentration of 0.01±0.00 and 10.26±4.75 g_COD_ L^-1^, respectively, treating 100% sewage sludge and 40% food waste with 60% sewage sludge, was measured in stage-II of two-stage systems with carriers (Figure 4b).

### 2.6 Electrical conductivity

During the first 60 days of sewage sludge mono-digestion, an electrical conductivity, which is an estimation of the overall salinity, of 4.7±1.6 and 4.9±1.4 mS cm^-1^ was observed in the single-stage systems without and with carriers, respectively. Maximum conductivity values of 25.8±0.3 mS cm^-1^ and 26.6±0.8 mS cm^-1^ were measured on day 195 in the single-stage systems without and with carriers, respectively at a maximum OLR of 3.8 g_VS_ L^-1^ d^-1^ treating a 40% food waste and 60% sewage sludge co-feedstock. Immediately after, the conductivity declined to average values of 19.4±0.3 and 21.3±0.3 mS cm^-1^ in the reactor without and with carriers, respectively treating 40% food waste and 60% sewage sludge co-feedstock on day 203 (Figure 5a).

**Figure 5:**
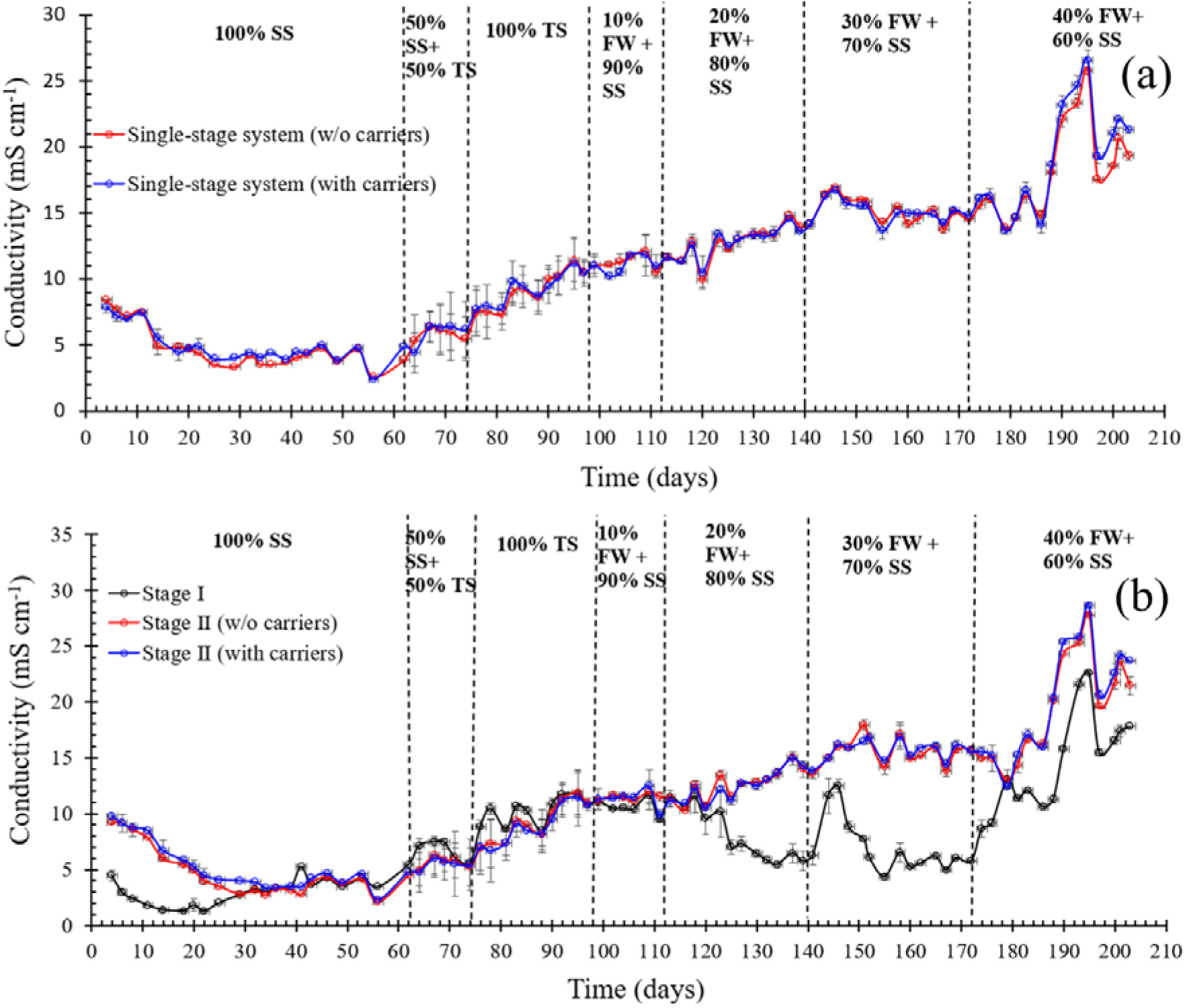
Conductivity in (a) the single-stage reactor without carriers (red line) and with carriers (blue line), and in (b) stage-I (black line), stage-II without carriers (red line) and stage-II with carriers (blue line) of the two-stage system. Data is expressed as mean ± standard deviation of biological duplicates (n=2). SS = Sewage Sludge, TS = Thickened Sludge, and FW = Food Waste.

In the two-stage systems, a higher fluctuation in the conductivity values was observed in stage-I compared to stage-II. At the start of the experimental period, the conductivity in stage-I was lower than in stage-II, with values lower than 5 mS cm^-1^. From day 63 on, the conductivity steadily increased in all reactors until day 118, after the introduction of 20% food waste into the co-feedstock (Figure 5b). The stage-I reactors exhibited a drop in the conductivity before a sharp increase to a local maximum of 12.5±0.6 mS cm^-1^ on day 146, while the stage-II reactors continued with the gradual increasing trend until day 146. When the food waste proportion was increased to 40%, the conductivity increased rapidly to 22.6±0.1 mS cm^-1^ in stage-I of two-stage system. The conductivity reached maximum values of 27.8±0.1 and 28.6±0.3 mS cm^-1^ in stage-II reactors of two-stage system without and with carriers, respectively, on day 195. Afterwards, conductivity values decreased to 17.8±0.0 mS cm^-1^ in stage-I, and 21.4±0.8 and 23.6±0.1 mS cm^-1^ in stage-II reactors without and with carriers, respectively at the end of the experimental period. Overall, the presence of carriers did not seem to have a distinct impact on the conductivity in either the single- or two-stage systems.

### 2.7 Volatile solids removal

The single-stage reactors with carriers outperformed the reactors without carriers in terms of VS reduction when treating different (co-)feedstocks. A VS reduction of 26.1±2.3% was observed in single-stage reactor with carriers treating 100% sewage sludge, which was twice as high in comparison with a VS reduction of 12.1±3.2% in single-stage reactor without carriers. With an increase in OLR by increasing the food waste proportion to 40% in the co-feedstock, an increase in VS reduction was noticed in the single-stage reactors with carriers, compared to reactors without carriers. The single-stage reactors without carriers showed a decreased VS removal efficiency from 77.2±2.5% at a 30% food waste and 70% sewage sludge co-feedstock to 65.5±11.6% treating 40% food waste and 60% sewage sludge (Figure 6a). A slight increase in VS reduction was observed in the single-stage with carriers from 78.8±3.7% at a 30% food waste and 70% sewage sludge co-feedstock to 81.4±5.9% when food waste proportion increased to 40%, which showed overall better performance at a maximum OLR of 3.8 g_VS_ L^-1^ d^-1^ (Figure 6a).

**Figure 6:**
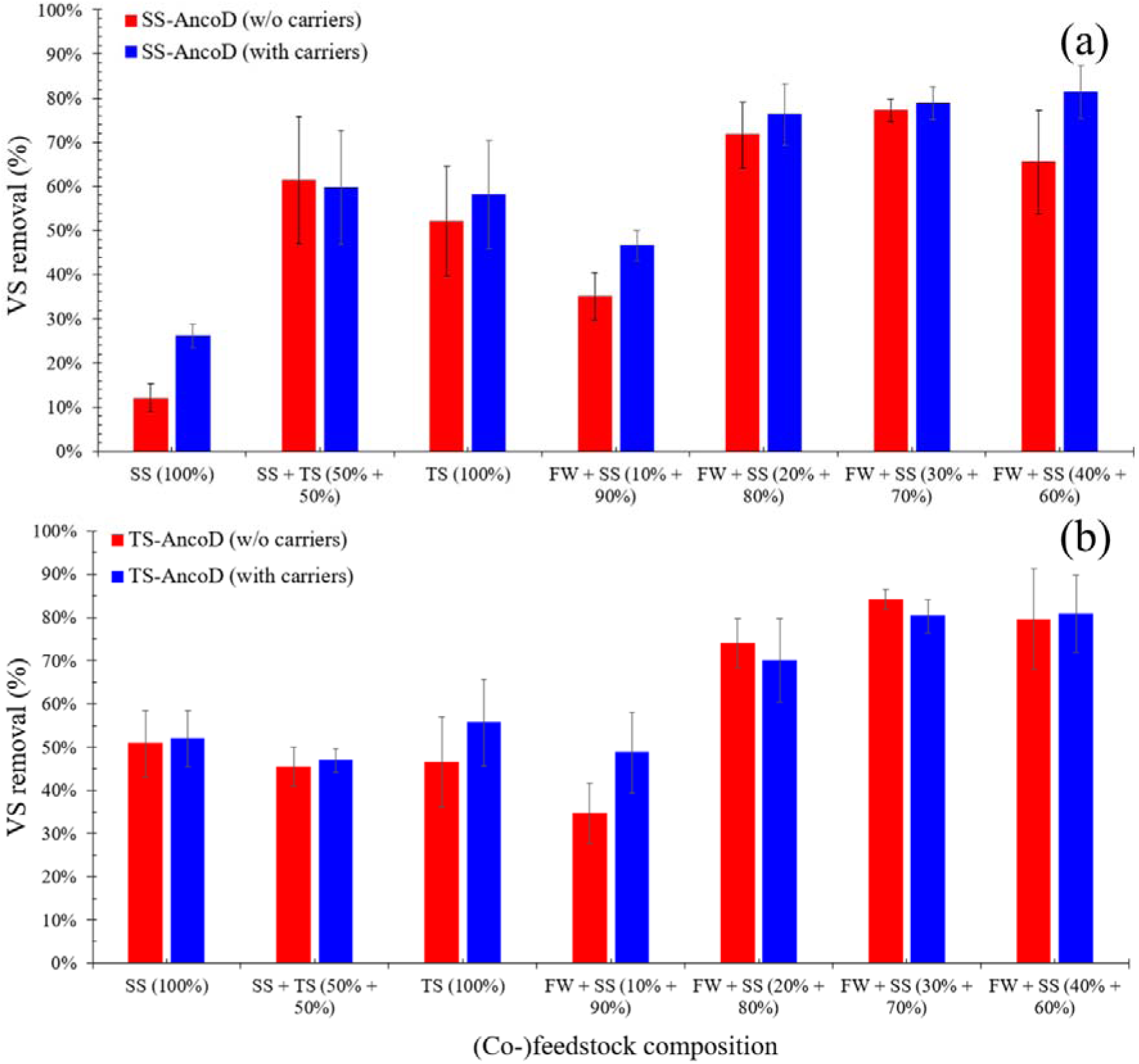
The volatile solids (VS) removal efficiency (%) in (a) the single-stage system without carriers (red block) and with carriers (blue block), and (b) the two-stage system without carriers (red block) and with carriers (blue block). Data is expressed as mean ± standard deviation of biological duplicates (n=2). SS = Sewage Sludge; TS = Thickened Sludge; FW = Food Waste, and VS = Volatile Solids.

An improved VS removal efficiency from 50.9±7.7% at a feedstock of 100% sewage sludge to 84.1±2.3% treating 30% food waste and 70% sewage sludge was observed in the two-stage reactors without carriers. When the food waste fraction was further increased to 40% in co-feedstock, a reduction in VS removal efficiency to 79.6±11.7% was noticed in the same reactors. The two-stage reactor with carriers showed an improved VS removal efficiency of 48.7±9.4% in comparison with the two-stage reactors without carriers of 34.7±7.0% treating co-feedstock of 10% food waste and 90% sewage sludge. A maximum VS removal efficiency of 80.9±9.0% was found in two-stage system with carriers co-digesting 40% food waste and 60% sewage sludge (Figure 6b).

## 3. Discussion

Co-digestion of food waste and sewage sludge resulted in more biogas than mono-digestion of sewage sludge in both single- and the two-stage systems. A higher biogas production rate was observed in the two-stage system, as both stages *i.e.*, stage-I and stage-II, produced biogas, compared to the single-stage system. System with carriers demonstrated a better system stability by producing less residual VFA, and a nominal increase in biogas production rate was also observed compared to a system without carriers.

### 3.1 Two-stage co-digestion of sewage sludge and food waste greatly outperformed the one-stage approach

The two-stage system exhibited overall higher biogas production rates than the single-stage system during the experimental period of 205 days, related to the fact that biogas was produced in both stages, *i.e.*, stage-I and stage-II of two-stage system. Schievano et al., 2023 [22] reported a very high biogas production rate of 3510±580 mL L^-1^ d^-1^ at HRT of 3 days, which corresponded to an OLR of 8.9±1.0 g_VS_ L^-1^ d^-1^ in stage-I of the two-stage system. It can be attributed to the short HRT that favours the presence of hydrolytic and acidogenic bacteria [41] which can result in an accumulation of VFA within stage-I of the two-stage system [42]. Paranjpe et al., 2023 [8] compared the biogas production between single- and two-stage systems treating food waste and sewage sludge, and achieved a higher biogas production rate of 670 mL L^-1^ d^-1^ in the two-stage and 451 mL L^-1^ d^-1^ in the single-stage system at 60% food waste and 40% sewage sludge under mesophilic conditions. In another study, Parra-Orobio et al., 2021 [43] reported 40% higher biogas production and 20% more methane content in a two-stage system treating the same co-feedstock.

Methane production in stage-I was most likely due to the presence and activity of hydrogenotrophic methanogenic archaea, related to the low HRT [44–46]. These hydrogenotrophic methanogenic archaea are more tolerant to a lower pH than acetoclastic methanogens, whose activity is generally inhibited at a pH below 6.2 [46–49]. Hydrogenotrophic methanogens also have a much lower minimum doubling time (6 to 12 hours), compared to acetoclastic methanogens (48 to 72 hours) [50–52]. This higher growth rate enables them to prevail at the shorter HRT values in stage-I, as they are less susceptible to washout [53]. The higher biogas production rate in stage-II of the two-stage system could be attributed to higher HRT, which enables slow growing acetoclastic methanogenic archaea to be involved in methanogenesis, in contrast to stage-I [54–56].

The methane yield obtained in the two-stage system was consistently higher than in the single-stage system throughout the experimental period, especially during the feeding regimes with 20% and 30% food waste. Kim et al., 2003 [57] reported a maximum methane yield of 215 mL CH_4_ g^-1^_VS_ when co-digesting food waste and sewage sludge at a ratio of 1:4, which was 85% higher than the methane yield obtained by mono-digestion of food waste. This methane yield value is lower than the 420±41 mL CH_4_ g^-1^_VS_ that was obtained in the single-stage system and 490±24 mL CH_4_ g^-1^_VS_ obtained in the two-stage system in our study treating the same co-feedstock. In another study, Liu et al., 2019 [58] reported a maximum methane yield of 554 mL CH_4_ g^-1^_VS_ at an OLR of 4.4 g_VS_ L^-1^ d^-1^. Other studies showed a 20-40% higher methane yield in two-stage systems compared to single-stage systems [59] and higher process stability [60,61], as summarized in Table 3.

**Table 3.** Maximum methane yields at different ratios of food waste and sewage sludge co-feedstock comparing literature data with our results.

| Sr. # | Co-feedstock composition | Methane Yield<br>(mL CH<sub>4</sub> g<sup>-1</sup> vs) | References |
| --- | --- | --- | --- |
| 1. | Food Waste: Sewage Sludge (20%: 80%) | 215-510 | (Kim et al., 2003; This study) |
| 2. | Food Waste: Sewage Sludge (25%: 75%) | 378 | (Varsha et al., 2022) |
| 3. | Food Waste: Sewage Sludge (30%: 70%) | 425 | This study |
| 4. | Food Waste: Sewage Sludge (40%: 60%) | 190 | This study |
| 5. | Food Waste: Sewage Sludge (50%: 50%) | 370-554 | (Heo et al., 2003; Y. Li et al., 2017; Varsha et al., 2022; Borowski, 2015; Liu et al., 2019) |
| 6. | Food Waste: Sewage Sludge (75%: 25%) | 333-720 | (Varsha et al., 2022; Liu et al., 2012) |
| 7. | Food Waste: Sewage Sludge (80%: 20%) | 500 | (Zhang et al., 2022) |

In our study, VS removal also showed a better performance in the two-stage system than in the single-stage system treating food waste and sewage sludge at different OLR values, as also reported earlier [62,63]. Another study showed that the VS removal efficiency remained below 20% when the OLR was increased from 3 to 6 g_VS_ L^-1^ d^-1^ in single-stage configuration, and reached up to 60% at an OLR of 7.3 g_VS_ L^-1^ d^-1^ in a two-stage configuration system [64]. Improved VS removal efficiencies of 5, 7, 8 and 48% have been reported in two-stage system having a total HRT of 20, 20, 30 and 12 days, respectively, in comparison with single-stage control reactors treating food waste and sewage sludge co-feedstock [65–67]. Two-stage systems reduce the risk for overloading, due to stage separation, and achieve an improved VS removal efficiency at higher OLR [68,69], which was also observed in our study.

### 3.2 A two-stage system showed higher process stability during co-digestion of sewage sludge and food waste than a single-stage system

The observed changes in pH and residual VFA concentrations reflected system stability, as both parameters are considered important indicators of anaerobic (co-)digestion stability [70]. In this study, a rapid process failure was observed at an increased OLR in both the single- and two-stage systems, primarily due to VFA accumulation [71–74] and salinity built-up [73–76], which negatively impacted the biogas production rate and methane yield. A sharp decline in pH has often been observed at an increasing OLR, due to accumulation of VFA, ultimately resulting in system failure [64,77–79]. In this study, a similar trend was also observed in the single-stage systems, wherein a substantial drop in pH was noticed, due to a rapid VFA accumulation, ultimately, resulting in ceased biogas production. Higher VFA accumulation is attributed to the lower methanogenic efficiency to assimilate VFA. Methanogens are more sensitive to salinity build-up than other fermentative microbes, and their growth can be inhibited and cell death can occur through dehydration [80,81]. In our study, a methane yield less than 50 mL CH_4_ g^-1^ was measured at a maximum conductivity of 18.1±0.3 mS cm^-1^ in the single-stage systems. Methanogens often lack robust mechanism for osmolytes synthesis (like ectoine), compared to other fermentative microbes [82–84] to tackle high salinity issue within the reactors, thus, leading to system failure. In the two-stage system, a stable pH of 7 was observed when the OLR was increased from 5 to 7 g_VS_ L^-1^ d^-1^ and drastically dropped to pH of 5.3 at an OLR of 10 g_VS_ L^-1^ d^-1^ [85]. In the present study, both the stages *i.e.*, stage-I and stage-II of the two-stage system followed the same trend. A sharp decline in pH was noticed much earlier in stage-I (day 112) compared to stage-II (day 178), which was due to VFA accumulation and high salinity built-up in stage-I of the two-stage system. A sudden drop in pH was observed, due to rapid acidification, which caused more VFA accumulation in stage-I compared to the stage-II reactors, reflecting an imbalance between VFA producing bacteria and consuming methanogenic archaea [86]. In stage-I of the two-stage system, biogas production was also ceased at much higher OLR of 15.2 g_VS_ L^-1^ d^-1^, and a conductivity of 22.6±0.1 mS cm^-1^ further inhibited methanogenic activity, as reported earlier [40,79,87]. Stage-II reactors maintained a neutral pH range much longer (till day 186), due to lesser VFA accumulation, compared to stage-I reactors wherein pH dropped to below 6 (on day 123). Overall, better system performance of two-stage system was observed in this study, due to efficient conversion of VFA to methane, reducing VFA accumulation and resulting in a stable pH at higher food waste amounts in the co-feedstock than in single-stage systems [88–96].

### 3.3 Carriers have a limited but positive contribution to biogas production

The use of carriers had an overall positive effect on the anaerobic digestion process stability. Higher VFA values were observed in the digesters without carriers than in those with carriers in both single- and two-stage systems. A more pronounced pH decrease in reactors without carriers ultimately resulting in process failure [97]. The addition of carriers in anaerobic digesters provides an additional surface area for biofilms to grow, and this biofilm structure allows slow growing methanogens to grow well in localized micro-environments with a more stable pH than the surrounding liquid phase in the reactor. Similar results were reported by Zhao et al., 2019 [98] when co-digesting sewage sludge and kitchen waste in ratios of 1:1, 1:2 and 1:3, using bentonite as a carrier material. The digesters with bentonite showed a pH range of 6.5 to 7.5, due to its high swelling property, which resulted in an increase surface area for biofilm development and improved growth of the methanogenic community. Similarly, in our study, a more stable pH and higher biogas production rates were observed in reactors with carriers. Biofilm formation on carriers most likely enabled higher microbial density and enhanced microbial synergy, and lead to efficient volatilization of solids [14,99–102]. Different studies have also showed an overall improved VS reduction in reactors with carriers enabling microbes to efficiently convert VFA and other intermediates into biogas [38,90,96,103,107–110]. Hamrouni and Cheikh 2021 [111] reported a maximum VS reduction of 86% in reactors with carriers at an OLR of 2.5 g_VS_ L^-1^ d^-1^, for a co-feedstock containing of 25% food waste and 75% sewage sludge. In other study, a maximum VS removal rate of 70% was achieved at an OLR of 6 g_VS_ L^-1^ d^-1^ in reactors with carriers treating same co-feedstock, as VS removal depends on easily available degradable matter present in co-feedstock [6,112]. A similar trend was also observed in our study, and the possible reasons could be the increased biofilm formation, due to the attachment of microbes on the carriers. Biofilms are dense microbial communities that are more efficient in degrading the organic matter than free floating microbes in reactors without carriers [107,113,114]. Biofilms formed on carriers also provide a shield to sensitive microbes against inhibitory substances, and allow slow-growing methanogens to function more efficiently [115–118]. Faisal et al., 2022 [97] studied the effect of low-density polyethylene (LDPET) and high-density polyethylene (HDPET) carriers, and reported an improved biogas production by 13% and 18% in reactors with LDPET and HDPET carriers, respectively, in comparison with a reactor without carriers. The VS removal efficiency was 41% higher in digester with HDPET carriers than in those without carriers. In another study, Liu et al., 2017 [119] investigated four types of fibrous biofilm carriers (polypropylene, polyester, polyamide, and polyurethane), and reported 10-45% higher biogas production and a 40-70% improved VS reduction in comparison with reactors without carriers. In our study, a maximum 25% increase in biogas production was observed and VS reduction was improved by 33% in two-stage system with carriers compared to reactors without carriers. Hence, systems with carriers often result in better system stability in comparison with those systems without carriers.

## 4. Conclusions

Co-digestion of sewage sludge with food waste resulted in an improved biogas production and VS reduction in comparison with mono-digestion of sewage sludge in both single- and two-stage systems. The use of carriers enhanced process stability in both single- and two-stage systems, but did not improve biogas production and VS reduction substantially. Stable process performance was observed until the food waste fraction reached 30% in the co-feedstock in both single- and two-stage reactors. A further increase in food waste to 40% resulted in a sharp decline in biogas production, resulting in process failure in both the single- and two-stage systems. Overall, the combination of 30% food waste and 70% sewage sludge co-feedstock performed best in terms of system performance and stability.

## CRediT author statement

**Muhammad Ali:** Conceptualization, Data Curation, Funding acquisition, Investigation, Methodology, Writing – Original Draft. **Carolina Burgos Pena:** Methodology, Formal Analysis, Investigation. **Jo De Vrieze:** Conceptualization, Data Curation, Funding acquisition, Resources, Supervision, Writing – Review & Editing.

## Acknowledgments

The authors would like to thank Aquafin NV for providing the inoculum and sewage sludge. Muhammad Ali received support from the Higher Education Commission (HEC), Pakistan under the scheme titled “Overseas Scholarship for PhD in selected fields Phase-III, Batch-3 with reference 2(3)/HRD/OSS-III/2022/HEC/583”. Jo De Vrieze acknowledges support from the Special Research Fund of Ghent University (BOF) with reference BOF/STA/202109/007.

## References

[1] S. Shaddel, H. Bakhtiary-Davijany, C. Kabbe, F. Dadgar, and S. W. Østerhus, “Sustainable Sewage Sludge Management: From Current Practices to Emerging Nutrient Recovery Technologies,” Sustainability, vol. 11, no. 12, Art. no. 12, Jan. 2019, doi: 10.3390/su11123435.

[2] M. M. Maghanaki, B. Ghobadian, G. Najafi, and R. J. Galogah, “Potential of biogas production in Iran,” Renew. Sustain. Energy Rev., vol. 28, pp. 702–714, Dec. 2013, doi: 10.1016/j.rser.2013.08.021.

[3] F. Baldi, I. Pecorini, and R. Iannelli, “Comparison of single-stage and two-stage anaerobic co-digestion of food waste and activated sludge for hydrogen and methane production,” Renew. Energy, vol. 143, pp. 1755–1765, Dec. 2019, doi: 10.1016/j.renene.2019.05.122.

[4] V. Khanh Nguyen et al., “Review on pretreatment techniques to improve anaerobic digestion of sewage sludge,” Fuel, vol. 285, p. 119105, Feb. 2021, doi: 10.1016/j.fuel.2020.119105.

[5] C. Aragon-Briceño et al., “Integration of hydrothermal carbonization treatment for water and energy recovery from organic fraction of municipal solid waste digestate,” Renew. Energy, vol. 184, pp. 577–591, Jan. 2022, doi: 10.1016/j.renene.2021.11.106.

[6] X. Dai, N. Duan, B. Dong, and L. Dai, “High-solids anaerobic co-digestion of sewage sludge and food waste in comparison with mono digestions: Stability and performance,” Waste Manag., vol. 33, no. 2, pp. 308–316, Feb. 2013, doi: 10.1016/j.wasman.2012.10.018.

[7] B. Karolinczak, J. Walczak, M. Bogacka, and M. Zubrowska-Sudol, “Life Cycle Assessment of sewage sludge mono-digestion and co-digestion with the organic fraction of municipal solid waste at a wastewater treatment plant,” Sci. Total Environ., vol. 907, p. 167801, Jan. 2024, doi: 10.1016/j.scitotenv.2023.167801.

[8] A. Paranjpe, S. Saxena, and P. Jain, “A Review on Performance Improvement of Anaerobic Digestion Using Co-Digestion of Food Waste and Sewage Sludge,” J. Environ. Manage., vol. 338, p. 117733, Jul. 2023, doi: 10.1016/j.jenvman.2023.117733.

[9] W. Tian et al., “Effects of hydrothermal pretreatment on the mono- and co-digestion of waste activated sludge and wheat straw,” Sci. Total Environ., vol. 732, p. 139312, Aug. 2020, doi: 10.1016/j.scitotenv.2020.139312.

[10] S. S. V. Varsha, A. F. Soomro, Z. T. Baig, A. K. Vuppaladadiyam, S. Murugavelh, and E. Antunes, “Methane production from anaerobic mono- and co-digestion of kitchen waste and sewage sludge: synergy study on cumulative methane production and biodegradability,” Biomass Convers. Biorefinery, vol. 12, no. 9, pp. 3911–3919, Sep. 2022, doi: 10.1007/s13399-020-00884-x.

[11] Y. Zhang, H. Li, C. Liu, and Y. Cheng, “Influencing mechanism of high solid concentration on anaerobic mono-digestion of sewage sludge without agitation,” Front. Environ. Sci. Eng., vol. 9, no. 6, pp. 1108–1116, Dec. 2015, doi: 10.1007/s11783-015-0806-x.

[12] R. Dalke, D. Demro, Y. Khalid, H. Wu, and M. Urgun-Demirtas, “Current status of anaerobic digestion of food waste in the United States,” Renew. Sustain. Energy Rev., vol. 151, p. 111554, Nov. 2021, doi: 10.1016/j.rser.2021.111554.

[13] P. Pagliaccia, A. Gallipoli, A. Gianico, D. Montecchio, and C. M. Braguglia, “Single stage anaerobic bioconversion of food waste in mono and co-digestion with olive husks: Impact of thermal pretreatment on hydrogen and methane production,” Int. J. Hydrog. Energy, vol. 41, no. 2, pp. 905–915, Jan. 2016, doi: 10.1016/j.ijhydene.2015.10.061.

[14] H. Song, Y. Zhang, S. Kusch-Brandt, and C. J. Banks, “Comparison of Variable and Constant Loading for Mesophilic Food Waste Digestion in a Long-Term Experiment,” Energies, vol. 13, no. 5, Art. no. 5, Jan. 2020, doi: 10.3390/en13051279.

[15] H. Tong, Y.-W. Tong, and Y. H. Peng, “A comparative life cycle assessment on mono- and co-digestion of food waste and sewage sludge,” Energy Procedia, vol. 158, pp. 4166–4171, Feb. 2019, doi: 10.1016/j.egypro.2019.01.814.

[16] D. Bolzonella, P. Battistoni, C. Susini, and F. Cecchi, “Anaerobic codigestion of waste activated sludge and OFMSW: the experiences of Viareggio and Treviso plants (Italy),” Water Sci. Technol., vol. 53, no. 8, pp. 203–211, Apr. 2006, doi: 10.2166/wst.2006.251.

[17] T. El-Hasan, S. Aljbour, and H. Al-Hamaiedeh, “Anaerobic co-digestion of domestic sewage sludge and food waste for biogas production: A decentralized integrated management of sludge in Jordan,” Jul. 2021.

[18] D. Chakraborty, O. P. Karthikeyan, A. Selvam, and J. W. C. Wong, “Co-digestion of food waste and chemically enhanced primary treated sludge in a continuous stirred tank reactor,” Biomass Bioenergy, vol. 111, pp. 232–240, Apr. 2018, doi: 10.1016/j.biombioe.2017.06.002.

[19] R. Karki et al., “Anaerobic co-digestion: Current status and perspectives,” Bioresour. Technol., vol. 330, p. 125001, Jun. 2021, doi: 10.1016/j.biortech.2021.125001.

[20] M. Aminzadeh, M. J. Bardi, and H. Aminirad, “A new approach to enhance the conventional two-phase anaerobic co-digestion of food waste and sewage sludge,” J. Environ. Health Sci. Eng., vol. 19, no. 1, pp. 295–306, Jun. 2021, doi: 10.1007/s40201-020-00603-8.

[21] M. Aminzadeh, M. J. Bardi, and H. Aminirad, “A new approach to enhance the conventional two-phase anaerobic co-digestion of food waste and sewage sludge,” J. Environ. Health Sci. Eng., vol. 19, no. 1, pp. 295–306, Jun. 2021, doi: 10.1007/s40201-020-00603-8.

[22] Andrea Schievano et al., “Two-Stage vs Single-Stage Thermophilic Anaerobic Digestion: Comparison of Energy Production and Biodegradation Efficiencies | Environmental Science & Technology.” Accessed: May 08, 2023. [Online]. Available: https://pubs.acs.org/doi/full/10.1021/es301376n

[23] S. J. Grimberg, D. Hilderbrandt, M. Kinnunen, and S. Rogers, “Anaerobic digestion of food waste through the operation of a mesophilic two-phase pilot scale digester – Assessment of variable loadings on system performance,” Bioresour. Technol., vol. 178, pp. 226–229, Feb. 2015, doi: 10.1016/j.biortech.2014.09.001.

[24] I. Muha et al., “Do two-phase biogas plants separate anaerobic digestion phases? – A mathematical model for the distribution of anaerobic digestion phases among reactor stages,” Bioresour. Technol., vol. 132, pp. 414–418, Mar. 2013, doi: 10.1016/j.biortech.2012.12.031.

[25] G. De Gioannis, A. Muntoni, A. Polettini, R. Pomi, and D. Spiga, “Energy recovery from one- and two-stage anaerobic digestion of food waste,” Waste Manag., vol. 68, pp. 595–602, Oct. 2017, doi: 10.1016/j.wasman.2017.06.013.

[26] D.-Y. Lee, Y. Ebie, K.-Q. Xu, Y.-Y. Li, and Y. Inamori, “Continuous H2 and CH4 production from high-solid food waste in the two-stage thermophilic fermentation process with the recirculation of digester sludge,” Bioresour. Technol., vol. 101, no. 1, Supplement, pp. S42–S47, Jan. 2010, doi: 10.1016/j.biortech.2009.03.037.

[27] G. Luo, L. Xie, Q. Zhou, and I. Angelidaki, “Enhancement of bioenergy production from organic wastes by two-stage anaerobic hydrogen and methane production process,” Bioresour. Technol., vol. 102, no. 18, pp. 8700–8706, Sep. 2011, doi: 10.1016/j.biortech.2011.02.012.

[28] M. A. Voelklein, A. Jacob, R. O’ Shea, and J. D. Murphy, “Assessment of increasing loading rate on two-stage digestion of food waste,” Bioresour. Technol., vol. 202, pp. 172–180, Feb. 2016, doi: 10.1016/j.biortech.2015.12.001.

[29] B. Xiao et al., “Comparison of single-stage and two-stage thermophilic anaerobic digestion of food waste: Performance, energy balance and reaction process,” Energy Convers. Manag., vol. 156, pp. 215–223, Jan. 2018, doi: 10.1016/j.enconman.2017.10.092.

[30] G. Srisowmeya, M. Chakravarthy, and G. Nandhini Devi, “Critical considerations in two-stage anaerobic digestion of food waste – A review,” Renew. Sustain. Energy Rev., vol. 119, p. 109587, Mar. 2020, doi: 10.1016/j.rser.2019.109587.

[31] I. M. C. Lo and K. S. Woon, “Food waste collection and recycling for value-added products: potential applications and challenges in Hong Kong,” Environ. Sci. Pollut. Res., vol. 23, no. 8, pp. 7081–7091, Apr. 2016, doi: 10.1007/s11356-015-4235-y.

[32] H.-W. Kim, J.-Y. Nam, and H.-S. Shin, “A comparison study on the high-rate co-digestion of sewage sludge and food waste using a temperature-phased anaerobic sequencing batch reactor system,” Bioresour. Technol., vol. 102, no. 15, pp. 7272– 7279, Aug. 2011, doi: 10.1016/j.biortech.2011.04.088.

[33] X. Liu, R. Li, and M. Ji, “Effects of Two-Stage Operation on Stability and Efficiency in Co-Digestion of Food Waste and Waste Activated Sludge,” Energies, vol. 12, no. 14, Art. no. 14, Jan. 2019, doi: 10.3390/en12142748.

[34] J.-H. Park, S.-H. Lee, J.-J. Yoon, S.-H. Kim, and H.-D. Park, “Predominance of cluster I *Clostridium* in hydrogen fermentation of galactose seeded with various heat-treated anaerobic sludges,” Bioresour. Technol., vol. 157, pp. 98–106, Apr. 2014, doi: 10.1016/j.biortech.2014.01.081.

[35] W. Li, J. Guo, H. Cheng, W. Wang, and R. Dong, “Two-phase anaerobic digestion of municipal solid wastes enhanced by hydrothermal pretreatment: Viability, performance and microbial community evaluation,” Appl. Energy, vol. 189, pp. 613–622, Mar. 2017, doi: 10.1016/j.apenergy.2016.12.101.

[36] P. Cui, J. Ge, Y. Chen, Y. Zhao, S. Wang, and H. Su, “The Fe3O4 nanoparticles-modified mycelium pellet-based anaerobic granular sludge enhanced anaerobic digestion of food waste with high salinity and organic load,” Renew. Energy, vol. 185, pp. 376–385, Feb. 2022, doi: 10.1016/j.renene.2021.12.050.

[37] M. Jiang et al., “Mitigating membrane fouling in a high solid food waste thermophilic anaerobic membrane bioreactor by incorporating fixed bed bio-carriers,” Chemosphere, vol. 292, p. 133488, Apr. 2022, doi: 10.1016/j.chemosphere.2021.133488.

[38] B. Wu et al., “Enhancing Anaerobic Digestion of Food Waste by Combining Carriers and Microaeration: Performance and Potential Mechanisms,” ACS EST Eng., Aug. 2024, doi: 10.1021/acsestengg.4c00298.

[39] N. A. F. Zamrisham, S. Idrus, M. R. Harun, M. S. A. Razak, and K. Jaman, “Biogas production by integrating lava rock, red clay & ceramic bio ring as support carrier in treatment of landfill leachate with liquidised food waste,” Biochem. Eng. J., vol. 204, p. 109221, Apr. 2024, doi: 10.1016/j.bej.2024.109221.

[40] S. Wang, X. Hou, and H. Su, “Exploration of the relationship between biogas production and microbial community under high salinity conditions,” Sci. Rep., vol. 7, no. 1, p. 1149, Apr. 2017, doi: 10.1038/s41598-017-01298-y.

[41] “Evaluation of biogas production and pollutant removal efficiency of two-phase anaerobic digestion treating slaughterhouse effluent: Biofuels: Vol 14, No 9.” Accessed: Mar. 03, 2025. [Online]. Available: https://www.tandfonline.com/doi/abs/10.1080/17597269.2023.2185728

[42] Z. Yang et al., “Dual optimization in anaerobic digestion of rice straw: Effects HRT and OLR coupling on methane production in one-stage and two-stage systems,” J. Environ. Manage., vol. 370, p. 123041, Nov. 2024, doi: 10.1016/j.jenvman.2024.123041.

[43] “Physicochemical, microbiological characterization and phytotoxicity of digestates produced on single-stage and two-stage anaerobic digestion of food waste | Sustainable Environment Research.” Accessed: Mar. 04, 2025. [Online]. Available: https://link.springer.com/article/10.1186/s42834-021-00085-9

[44] “The effect of temperature and retention time on methane production and microbial community composition in staged anaerobic digesters fed with food waste | Biotechnology for Biofuels and Bioproducts.” Accessed: Mar. 03, 2025. [Online]. Available: https://link.springer.com/article/10.1186/s13068-017-0989-4

[45] T. Shimada et al., “Syntrophic acetate oxidation in two-phase (acid–methane) anaerobic digesters,” Water Sci. Technol., vol. 64, no. 9, pp. 1812–1820, Nov. 2011, doi: 10.2166/wst.2011.748.

[46] B. Demirel and P. Scherer, “Production of methane from sugar beet silage without manure addition by a single-stage anaerobic digestion process,” Biomass Bioenergy, vol. 32, no. 3, pp. 203–209, Mar. 2008, doi: 10.1016/j.biombioe.2007.09.011.

[47] M. Sun et al., “Effects of low pH conditions on decay of methanogenic biomass,” Water Res., vol. 179, p. 115883, Jul. 2020, doi: 10.1016/j.watres.2020.115883.

[48] C. Wang, Y. Li, and Y. Sun, “Acclimation of Acid-Tolerant Methanogenic Culture for Bioaugmentation: Strategy Comparison and Microbiome Succession,” ACS Omega, vol. 5, no. 11, pp. 6062–6068, Mar. 2020, doi: 10.1021/acsomega.9b03783.

[49] L. Zhang, X. Liu, K. Duddleston, and M. E. Hines, “The Effects of pH, Temperature, and Humic-Like Substances on Anaerobic Carbon Degradation and Methanogenesis in Ombrotrophic and Minerotrophic Alaskan Peatlands,” Aquat. Geochem., vol. 26, no. 3, pp. 221–244, Sep. 2020, doi: 10.1007/s10498-020-09372-0.

[50] D. Dzofou Ngoumelah et al., “A unified and simple medium for growing model methanogens,” Front. Microbiol., vol. 13, Jan. 2023, doi: 10.3389/fmicb.2022.1046260.

[51] Z. Lü and Y. Lu, “Methanocella conradii sp. nov., a Thermophilic, Obligate Hydrogenotrophic Methanogen, Isolated from Chinese Rice Field Soil,” PLOS ONE, vol. 7, no. 4, p. e35279, Apr. 2012, doi: 10.1371/journal.pone.0035279.

[52] Y. Song, W. Qiao, J. Zhang, and R. Dong, “Process Performance and Functional Microbial Community in the Anaerobic Digestion of Chicken Manure: A Review,” Energies, vol. 16, no. 12, Art. no. 12, Jan. 2023, doi: 10.3390/en16124675.

[53] Y.-X. Huang, J. Guo, C. Zhang, and Z. Hu, “Hydrogen production from the dissolution of nano zero valent iron and its effect on anaerobic digestion,” Water Res., vol. 88, pp. 475–480, Jan. 2016, doi: 10.1016/j.watres.2015.10.028.

[54] K. Ignatowicz, G. Filipczak, B. Dybek, and G. Wałowski, “Biogas Production Depending on the Substrate Used: A Review and Evaluation Study—European Examples,” Energies, vol. 16, no. 2, Art. no. 2, Jan. 2023, doi: 10.3390/en16020798.

[55] R. Nkuna, A. Roopnarain, C. Rashama, and R. Adeleke, “Insights into organic loading rates of anaerobic digestion for biogas production: a review,” Crit. Rev. Biotechnol., vol. 42, no. 4, pp. 487–507, May 2022, doi: 10.1080/07388551.2021.1942778.

[56] S. Yang et al., “Biogas production of food waste with *in-situ* sulfide control under high organic loading in two-stage anaerobic digestion process: Strategy and response of microbial community,” Bioresour. Technol., vol. 373, p. 128712, Apr. 2023, doi: 10.1016/j.biortech.2023.128712.

[57] J. Kim et al., “Effects of various pretreatments for enhanced anaerobic digestion with waste activated sludge,” J. Biosci. Bioeng., vol. 95, no. 3, pp. 271–275, Jan. 2003, doi: 10.1016/S1389-1723(03)80028-2.

[58] X. Liu et al., “Mechanistic insights into the effect of poly ferric sulfate on anaerobic digestion of waste activated sludge,” Water Res., vol. 189, p. 116645, Feb. 2021, doi: 10.1016/j.watres.2020.116645.

[59] A. J. Guneratnam et al., “Study of the performance of a thermophilic biological methanation system,” Bioresour. Technol., vol. 225, pp. 308–315, Feb. 2017, doi: 10.1016/j.biortech.2016.11.066.

[60] K. Rajendran, D. Mahapatra, A. V. Venkatraman, S. Muthuswamy, and A. Pugazhendhi, “Advancing anaerobic digestion through two-stage processes: Current developments and future trends,” Renew. Sustain. Energy Rev., vol. 123, p. 109746, May 2020, doi: 10.1016/j.rser.2020.109746.

[61] M. A. Voelklein, R. O’ Shea, A. Jacob, and J. D. Murphy, “Role of trace elements in single and two-stage digestion of food waste at high organic loading rates,” Energy, vol. 121, pp. 185–192, Feb. 2017, doi: 10.1016/j.energy.2017.01.009.

[62] J. A. Arzate et al., “Anaerobic Digestion Model (AM2) for the Description of Biogas Processes at Dynamic Feedstock Loading Rates,” Chem. Ing. Tech., vol. 89, no. 5, pp. 686–695, 2017, doi: 10.1002/cite.201600176.

[63] C. E. Gómez Camacho, B. Ruggeri, L. Mangialardi, M. Persico, and A. C. Luongo Malavé, “Continuous two-step anaerobic digestion (TSAD) of organic market waste: rationalising process parameters,” Int. J. Energy Environ. Eng., vol. 10, no. 4, pp. 413–427, Dec. 2019, doi: 10.1007/s40095-019-0312-1.

[64] B. A. Parra-Orobio, M. N. Cruz-Bournazou, and P. Torres-Lozada, “Single-Stage and Two-Stage Anaerobic Digestion of Food Waste: Effect of the Organic Loading Rate on the Methane Production and Volatile Fatty Acids,” Water. Air. Soil Pollut., vol. 232, no. 3, p. 105, Mar. 2021, doi: 10.1007/s11270-021-05064-9.

[65] Y. Maspolim, Y. Zhou, C. Guo, K. Xiao, and W. J. Ng, “Comparison of single-stage and two-phase anaerobic sludge digestion systems – Performance and microbial community dynamics,” Chemosphere, vol. 140, pp. 54–62, Dec. 2015, doi: 10.1016/j.chemosphere.2014.07.028.

[66] R. Nabaterega, V. Kumar, S. Khoei, and C. Eskicioglu, “A review on two-stage anaerobic digestion options for optimizing municipal wastewater sludge treatment process,” J. Environ. Chem. Eng., vol. 9, no. 4, p. 105502, Aug. 2021, doi: 10.1016/j.jece.2021.105502.

[67] J. D. Zahller, R. H. Bucher, J. F. Ferguson, and H. D. Stensel, “Performance and Stability of Two-Stage Anaerobic Digestion,” Water Environ. Res., vol. 79, no. 5, pp. 488–497, 2007, doi: 10.2175/106143006X123157.

[68] H. Chen et al., “Biohythane production and microbial characteristics of two alternating mesophilic and thermophilic two-stage anaerobic co-digesters fed with rice straw and pig manure,” Bioresour. Technol., vol. 320, p. 124303, Jan. 2021, doi: 10.1016/j.biortech.2020.124303.

[69] J. Rusín, K. Chamrádová, and P. Basinas, “Two-stage psychrophilic anaerobic digestion of food waste: Comparison to conventional single-stage mesophilic process,” Waste Manag., vol. 119, pp. 172–182, Jan. 2021, doi: 10.1016/j.wasman.2020.09.039.

[70] A. Ajayi-Banji and S. Rahman, “A review of process parameters influence in solid-state anaerobic digestion: Focus on performance stability thresholds,” Renew. Sustain. Energy Rev., vol. 167, p. 112756, Oct. 2022, doi: 10.1016/j.rser.2022.112756.

[71] P. P. Mathai et al., “Dynamic shifts within volatile fatty acid–degrading microbial communities indicate process imbalance in anaerobic digesters,” Appl. Microbiol. Biotechnol., vol. 104, no. 10, pp. 4563–4575, May 2020, doi: 10.1007/s00253-020-10552-9.

[72] K. Maurus, N. Kremmeter, S. Ahmed, and M. Kazda, “High-resolution monitoring of VFA dynamics reveals process failure and exponential decrease of biogas production,” Biomass Convers. Biorefinery, vol. 13, no. 12, pp. 10653–10663, Aug. 2023, doi: 10.1007/s13399-021-02043-2.

[73] A. L. Pera, M. Sellaro, M. Bianco, and G. Zanardi, “Effects of a temporary increase in OLR and a simultaneous decrease in HRT on dry anaerobic digestion of OFMSW,” Environ. Technol., vol. 43, no. 28, pp. 4463–4471, Dec. 2022, doi: 10.1080/09593330.2021.1952312.

[74] K. Xiao, C. Guo, Y. Zhou, Y. Maspolim, and W.-J. Ng, “Acetic acid effects on methanogens in the second stage of a two-stage anaerobic system,” Chemosphere, vol. 144, pp. 1498–1504, Feb. 2016, doi: 10.1016/j.chemosphere.2015.10.035.

[75] L. V. Duc, Y. Miyagawa, D. Inoue, and M. Ike, “Identification of key steps and associated microbial populations for efficient anaerobic digestion under high ammonium or salinity conditions,” Bioresour. Technol., vol. 360, p. 127571, Sep. 2022, doi: 10.1016/j.biortech.2022.127571.

[76] J. D. Muñoz Sierra, M. J. Oosterkamp, H. Spanjers, and J. B. van Lier, “Effects of large salinity fluctuations on an anaerobic membrane bioreactor treating phenolic wastewater,” Chem. Eng. J., vol. 417, p. 129263, Aug. 2021, doi: 10.1016/j.cej.2021.129263.

[77] M. Capodici, G. Mannina, and M. Torregrossa, “Waste activated sludge dewaterability: comparative evaluation of sludge derived from CAS and MBR systems,” Desalination Water Treat., vol. 57, no. 48–49, pp. 22917–22925, Oct. 2016, doi: 10.1080/19443994.2016.1180478.

[78] M. Peces, G. Pozo, K. Koch, J. Dosta, and S. Astals, “Exploring the potential of co-fermenting sewage sludge and lipids in a resource recovery scenario,” Bioresour. Technol., vol. 300, p. 122561, Mar. 2020, doi: 10.1016/j.biortech.2019.122561.

[79] R. Rajagopal, D. I. Massé, and G. Singh, “A critical review on inhibition of anaerobic digestion process by excess ammonia,” Bioresour. Technol., vol. 143, pp. 632–641, Sep. 2013, doi: 10.1016/j.biortech.2013.06.030.

[80] X. F. Li et al., “Study of Salt Effect on Semi-Continuous Anaerobic Digestion of Food Waste with Modified First-Order Model,” IOP Conf. Ser. Earth Environ. Sci., vol. 701, no. 1, p. 012032, Mar. 2021, doi: 10.1088/1755-1315/701/1/012032.

[81] Q. Meng, H. Liu, H. Zhang, S. Xu, E. Lichtfouse, and Y. Yun, “Anaerobic digestion and recycling of kitchen waste: a review,” Environ. Chem. Lett., vol. 20, no. 3, pp. 1745–1762, Jun. 2022, doi: 10.1007/s10311-022-01408-x.

[82] M. O. Fagbohungbe, I. C. Dodd, B. M. J. Herbert, H. Li, L. Ricketts, and K. T. Semple, “High solid anaerobic digestion: Operational challenges and possibilities,” *Environ*. Technol. Innov., vol. 4, pp. 268–284, Oct. 2015, doi: 10.1016/j.eti.2015.09.003.

[83] M. Gao et al., “Deep insights into the anaerobic co-digestion of waste activated sludge with concentrated leachate under different salinity stresses,” Sci. Total Environ., vol. 838, p. 155922, Sep. 2022, doi: 10.1016/j.scitotenv.2022.155922.

[84] X. He et al., “The impact of salinity on biomethane production and microbial community in the anaerobic digestion of food waste components,” Energy, vol. 294, p. 130736, May 2024, doi: 10.1016/j.energy.2024.130736.

[85] R. Ganesh, M. Torrijos, P. Sousbie, A. Lugardon, J. P. Steyer, and J. P. Delgenes, “Single-phase and two-phase anaerobic digestion of fruit and vegetable waste: Comparison of start-up, reactor stability and process performance,” Waste Manag., vol. 34, no. 5, pp. 875–885, May 2014, doi: 10.1016/j.wasman.2014.02.023.

[86] S. Aslanzadeh, K. Rajendran, and M. J. Taherzadeh, “A comparative study between single- and two-stage anaerobic digestion processes: Effects of organic loading rate and hydraulic retention time,” Int. Biodeterior. Biodegrad., vol. 95, pp. 181–188, Nov. 2014, doi: 10.1016/j.ibiod.2014.06.008.

[87] L. Appels, J. Baeyens, J. Degrève, and R. Dewil, “Principles and potential of the anaerobic digestion of waste-activated sludge,” Prog. Energy Combust. Sci., vol. 34, no. 6, pp. 755–781, Dec. 2008, doi: 10.1016/j.pecs.2008.06.002.

[88] F. Almomani, R. R. Bhosale, M. A. M. Khraisheh, and M. Shawaqfah, “Enhancement of biogas production from agricultural wastes via pre-treatment with advanced oxidation processes,” Fuel, vol. 253, pp. 964–974, Oct. 2019, doi: 10.1016/j.fuel.2019.05.057.

[89] J. De Vrieze, A. Devooght, D. Walraedt, and N. Boon, “Enrichment of Methanosaetaceae on carbon felt and biochar during anaerobic digestion of a potassium-rich molasses stream,” Appl. Microbiol. Biotechnol., vol. 100, no. 11, pp. 5177–5187, Jun. 2016, doi: 10.1007/s00253-016-7503-y.

[90] W. Gong, H. Liang, W. Li, and Z. Wang, “Selection and evaluation of biofilm carrier in anaerobic digestion treatment of cattle manure,” Energy, vol. 36, no. 5, pp. 3572–3578, May 2011, doi: 10.1016/j.energy.2011.03.068.

[91] J. N. Haigh, T. R. Dargaville, and P. D. Dalton, “Additive manufacturing with polypropylene microfibers,” Mater. Sci. Eng. C, vol. 77, pp. 883–887, Aug. 2017, doi: 10.1016/j.msec.2017.03.286.

[92] X. Huang, S. Yun, J. Zhu, T. Du, C. Zhang, and X. Li, “Mesophilic anaerobic co-digestion of aloe peel waste with dairy manure in the batch digester: Focusing on mixing ratios and digestate stability,” Bioresour. Technol., vol. 218, pp. 62–68, Oct. 2016, doi: 10.1016/j.biortech.2016.06.070.

[93] Y. Liu et al., “Effects of amino-modified biofilm carriers on biogas production in the anaerobic digestion of corn straw,” Environ. Technol., vol. 41, no. 21, pp. 2806–2816, Sep. 2020, doi: 10.1080/09593330.2019.1583290.

[94] N. Qureshi, B. A. Annous, T. C. Ezeji, P. Karcher, and I. S. Maddox, “Biofilm reactors for industrial bioconversion processes: employing potential of enhanced reaction rates,” Microb. Cell Factories, vol. 4, no. 1, p. 24, Aug. 2005, doi: 10.1186/1475-2859-4-24.

[95] W. Wu et al., “Improving methane production in cow dung and corn straw co-fermentation systems via enhanced degradation of cellulose by cabbage addition,” Sci. Rep., vol. 6, no. 1, p. 33628, Sep. 2016, doi: 10.1038/srep33628.

[96] A. Zainab, S. Meraj, and R. Liaquat, “Study on Natural Organic Materials as Biofilm Carriers for the Optimization of Anaerobic Digestion,” Waste Biomass Valorization, vol. 11, no. 6, pp. 2521–2531, Jun. 2020, doi: 10.1007/s12649-019-00628-7.

[97] S. Faisal et al., “Enhanced waste hot-pot oil (WHPO) anaerobic digestion for biomethane production: Mechanism and dynamics of fatty acids conversion,” Chemosphere, vol. 307, p. 135955, Nov. 2022, doi: 10.1016/j.chemosphere.2022.135955.

[98] “Understanding the mechanism of polybrominated diphenyl ethers reducing the anaerobic co-digestion efficiency of excess sludge and kitchen waste | Environmental Science and Pollution Research.” Accessed: Mar. 04, 2025. [Online]. Available: https://link.springer.com/article/10.1007/s11356-022-18795-x

[99] F. Bianco, M. Race, V. Forino, S. Pacheco-Ruiz, and E. R. Rene, “Chapter 4 - Bioreactors for wastewater to energy conversion: from pilot to full scale experiences,” in Waste Biorefinery, T. Bhaskar, S. Varjani, A. Pandey, and E. R. Rene, Eds., Elsevier, 2021, pp. 103–124. doi: 10.1016/B978-0-12-821879-2.00004-1.

[100] M. Pohl, K. Heeg, and J. Mumme, “Anaerobic digestion of wheat straw – Performance of continuous solid-state digestion,” Bioresour. Technol., vol. 146, pp. 408–415, Oct. 2013, doi: 10.1016/j.biortech.2013.07.101.

[101] I. Ramos and M. Fdz-Polanco, “The potential of oxygen to improve the stability of anaerobic reactors during unbalanced conditions: Results from a pilot-scale digester treating sewage sludge,” Bioresour. Technol., vol. 140, pp. 80–85, Jul. 2013, doi: 10.1016/j.biortech.2013.04.066.

[102] Y. Wei, Y. Gao, H. Yuan, Y. Chang, and X. Li, “Effects of organic loading rate and pretreatments on digestion performance of corn stover and chicken manure in completely stirred tank reactor (CSTR),” Sci. Total Environ., vol. 815, p. 152499, Apr. 2022, doi: 10.1016/j.scitotenv.2021.152499.

[103] J. Liu, T. Liu, S. Chen, H. Yu, Y. Zhang, and X. Quan, “Enhancing anaerobic digestion in anaerobic integrated floating fixed-film activated sludge (An-IFFAS) system using novel electron mediator suspended biofilm carriers,” Water Res., vol. 175, p. 115697, May 2020, doi: 10.1016/j.watres.2020.115697.

[104] S. Montalvo et al., “Application of natural zeolites in anaerobic digestion processes: A review,” Appl. Clay Sci., vol. 58, pp. 125–133, Apr. 2012, doi: 10.1016/j.clay.2012.01.013.

[105] M. Nabi, H. Liang, L. Cheng, W. Yang, and D. Gao, “A comprehensive review on the use of conductive materials to improve anaerobic digestion: Focusing on landfill leachate treatment,” J. Environ. Manage., vol. 309, p. 114540, May 2022, doi: 10.1016/j.jenvman.2022.114540.

[106] S. Weiß et al., “Activated zeolite—suitable carriers for microorganisms in anaerobic digestion processes?,” Appl. Microbiol. Biotechnol., vol. 97, no. 7, pp. 3225–3238, Apr. 2013, doi: 10.1007/s00253-013-4691-6.

[107] K. Ding et al., “Study on synergistic effect of carrier combined with micro-aeration on anaerobic digestion of food waste,” Chem. Eng. J., vol. 498, p. 155731, Oct. 2024, doi: 10.1016/j.cej.2024.155731.

[108] L. Feng et al., “Mechanisms, performance, and the impact on microbial structure of direct interspecies electron transfer for enhancing anaerobic digestion-A review,” Sci. Total Environ., vol. 862, p. 160813, Mar. 2023, doi: 10.1016/j.scitotenv.2022.160813.

[109] Y. Zhang, C. Li, Z. Yuan, R. Wang, I. Angelidaki, and G. Zhu, “Syntrophy mechanism, microbial population, and process optimization for volatile fatty acids metabolism in anaerobic digestion,” Chem. Eng. J., vol. 452, p. 139137, Jan. 2023, doi: 10.1016/j.cej.2022.139137.

[110] D. Zhao et al., “Mitigation of acidogenic product inhibition and elevated mass transfer by biochar during anaerobic digestion of food waste,” Bioresour. Technol., vol. 338, p. 125531, Oct. 2021, doi: 10.1016/j.biortech.2021.125531.

[111] Y. M. B. Hamrouni and R. B. Cheikh, “Enhancing the energetic potential of Mediterranean food waste by anaerobic co-digestion with sewage sludge,” Environ. Prog. Sustain. Energy, vol. 40, no. 2, p. e13512, 2021, doi: 10.1002/ep.13512.

[112] C. Gou, Z. Yang, J. Huang, H. Wang, H. Xu, and L. Wang, “Effects of temperature and organic loading rate on the performance and microbial community of anaerobic co-digestion of waste activated sludge and food waste,” Chemosphere, vol. 105, pp. 146– 151, Jun. 2014, doi: 10.1016/j.chemosphere.2014.01.018.

[113] R. D. A. Cayetano et al., “Biofilm formation as a method of improved treatment during anaerobic digestion of organic matter for biogas recovery,” Bioresour. Technol., vol. 344, p. 126309, Jan. 2022, doi: 10.1016/j.biortech.2021.126309.

[114] Y. Qian, Y. Guo, J. Shen, Y. Qin, and Y.-Y. Li, “Biofilm growth characterization and treatment performance in a single stage partial nitritation/anammox process with a biofilm carrier,” Water Res., vol. 217, p. 118437, Jun. 2022, doi: 10.1016/j.watres.2022.118437.

[115] S. Arif, R. Liaquat, and M. Adil, “Applications of materials as additives in anaerobic digestion technology,” Renew. Sustain. Energy Rev., vol. 97, pp. 354–366, Dec. 2018, doi: 10.1016/j.rser.2018.08.039.

[116] S. Faisal, E.-S. Salama, S. H. A. Hassan, B.-H. Jeon, and X. Li, “Biomethane enhancement via plastic carriers in anaerobic co-digestion of agricultural wastes,” Biomass Convers. Biorefinery, vol. 12, no. 7, pp. 2553–2565, Jul. 2022, doi: 10.1007/s13399-020-00779-x.

[117] A. H. Y. Fung et al., “Exploring the optimization of aerobic food waste digestion efficiency through the engineering of functional biofilm Bio-carriers,” Bioresour. Technol., vol. 341, p. 125869, Dec. 2021, doi: 10.1016/j.biortech.2021.125869.

[118] A. H. Jagaba et al., “A systematic literature review of biocarriers: Central elements for biofilm formation, organic and nutrients removal in sequencing batch biofilm reactor,” J. Water Process Eng., vol. 42, p. 102178, Aug. 2021, doi: 10.1016/j.jwpe.2021.102178.

[119] T. Liu, X. Zhou, Z. Li, X. Wang, and J. Sun, “Effects of liquid digestate pretreatment on biogas production for anaerobic digestion of wheat straw,” Bioresour. Technol., vol. 280, pp. 345–351, May 2019, doi: 10.1016/j.biortech.2019.01.147.

