## Supplementary file 1 for "Anaerobic co-digestion of sewage sludge and food waste: staging and carriers enhance system performance and process stability"

**Supplementary Data:**

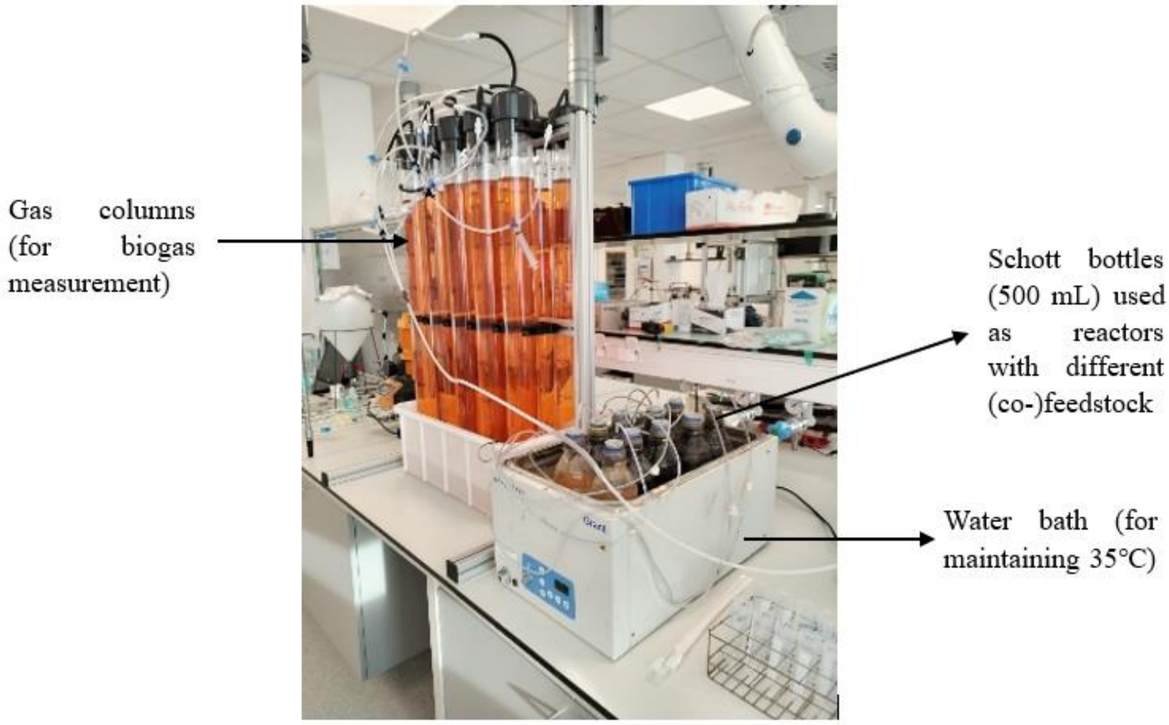


**Figure S1:** Actual lab-scale set-up of anaerobic (co-)digestion system

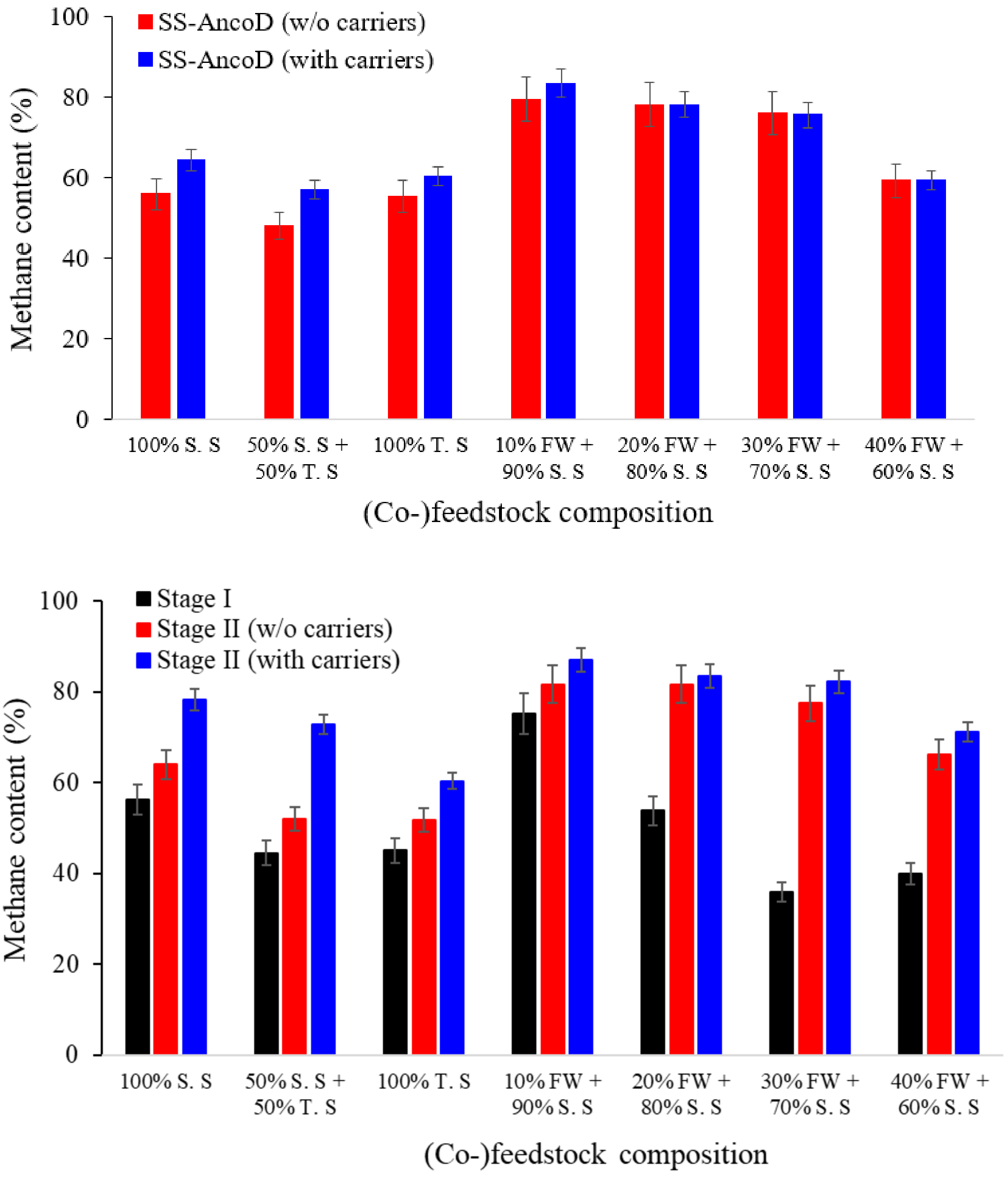


**Figure S2:** Methane content (%) in (a) the single-stage anaerobic co-digestion (SS-AncoD) reactor without carriers (red block) and with carriers (blue block), and (b) stage-I (black block), stage-II without carriers (red block) and stage-II with carriers (blue block) of the two-stage system. Data is expressed as mean ± standard deviation of biological duplicates (n=2).
